# Sparsity of higher-order landscape interactions enables learning and prediction for microbiomes

**DOI:** 10.1101/2023.04.12.536602

**Authors:** Shreya Arya, Ashish B. George, James P. O’Dwyer

## Abstract

Microbiome engineering offers the potential to leverage microbial communities to improve outcomes in human health, agriculture, and climate. To translate this potential into reality, it is crucial to reliably predict community composition and function. But a brute force approach to cataloguing community function is hindered by the combinatorial explosion in the number of ways we can combine microbial species. An alternative is to parameterize microbial community outcomes using simplified, mechanistic models, and then extrapolate these models beyond where we have sampled. But these approaches remain data-hungry, as well as requiring an *a priori* specification of what kinds of mechanism are included and which are omitted. Here, we resolve both issues by introducing a new, mechanism-agnostic approach to predicting microbial community compositions and functions using limited data. The critical step is the discovery of a sparse representation of the community landscape. We then leverage this sparsity to predict community compositions and functions, drawing from techniques in compressive sensing. We validate this approach on *in silico* community data, generated from a theoretical model. By sampling just ∼ 1% of all possible communities, we accurately predict community compositions out of sample. We then demonstrate the real-world application of our approach by applying it to four experimental datasets, and showing that we can recover interpretable, accurate predictions on composition and community function from highly limited data.

## Introduction

Our planet is host to a multitude of microbial communities, also known as microbiomes, which perform an enormous range of functions in shaping biogeochemical processes, agricultural productivity, and animal and human health [1–3]. In recent years, there have been concerted efforts to modify such communities in order to alter plant, animal, human, and environmental health for the better [4–14]. The complex interspecific interactions present in real communities lead to diverse steady-state communities, but these interactions also mean that the final composition and function of a microbiome may be very different from its initial composition. Moreover, for large numbers of potential taxa to include in a community, there is a combinatorially-large number of distinct initial compositions. Putting these two issues together makes a brute-force approach to cataloging potential microbiomes impossible: with as few as ten species from which to build an initial community, 2^10^, or around 1000 experiments would be needed to survey the full range of possible outcomes arising as the dynamics of communities play out. With 100 potential initial species, that number becomes 10^30^ [15].

Building mechanistic models of microbial interspecific interactions has the potential to alleviate this issue. If we can write down and parameterize a model that accurately represents interactions, but using only a limited amount of experimental data, we might be able to use such a model to predict outcomes beyond those experiments. A longstanding approach to modeling interspecific interactions, inspired by Robert May’s seminal work [16] on complex systems, has been to use pairwise interactions among species, with the strength of this pairwise dependence encoded in a community interaction matrix [17, 18]. There is certainly no conceptual obstacle to pairwise interactions providing a potential description of microbial communities—theoretical work demonstrates that pairwise interactions can lead to diverse, realistic communities [19–26]. But there is also a recent realization that microbial communities may be infused with higher-order interspecific interactions, where the influence of one species on another is dependent on the presence or absence of a third, fourth, or fifth species [27–31]. Even for pairwise interspecific interactions, having *S*^2^ parameters to fit means that mechanistic models could just bring us back to a similar experimental load of the combinatorial problem we started with [18, 27, 32, 33]. Higher-order interspecific interactions would then only make this problem of fitting parameters even more challenging. Finally, deep learning has been deployed to address this question [32, 34], but at the expense of making results and predictions challenging to interpret.

Developing a method that requires only limited experimental data, is agnostic to any particular ecological model, and provides ecologically-interpretable, accurate results, would add an important tool to the microbial ecologist’s toolbox. Here, we will address this need, introducing an approach that reveals a previously-hidden sparsity in the landscape describing the map from initial composition to final community states, for both simulated and experimental microbial communities. The sparsity of this landscape means that the outcomes of microbial community assembly are highly constrained, in effect containing much less information than the full combinatorial set would suggest. We then leverage this sparsity, using algorithms from compressive sensing to accurately recover the sparse representation. Thus, we predict the late-time outcomes for microbial community composition, from highly limited input data, in a way that is readily interpretable in terms of the sparsity of landscape interactions.

### Framework

#### The challenge of predicting microbial community composition

We first explicitly state the problem to address: given a set of *S* species to draw from, there are 2^*S*^ − 1 possible combinations, or seed communities, that can be formed, based on the initial presence-absence of species. This initial condition naturally does not capture the full subsequent behavior of the community, which could be extremely complex [35–38]. Here, we will make a simplifying assumption that, from a given initial condition, all species will approach an equilibrium at late times, such that each species ends up with a relative abundance that only depends on the seed community composition. This excludes more general dynamics, including chaos [39], limit cycles [40], or priority effects [41]. We justify this assumption by noting, as in [15], that this simple behavior is what has been frequently observed in experimental communities. Furthermore, this behavior is predicted by many mechanistic models (see Table 1 in [15]). More complex behavior can be observed in ecological communities, demonstrated both theoretically in terms of multiple attractors in Lotka-Volterra models [40], and experimentally, where for some systems either dynamical attractors or a single stable equilibrium can exist, dependent on the parameter regime [42]. But making progress in the case of systems with a unique steady-state may be a first step towards tackling these more general cases. Even with this assumption of a unique steady-state, if there are any interspecific interactions at all, the abundance of a species at steady state will in general depend in a complex way on which other species are present.

**Table 1.**
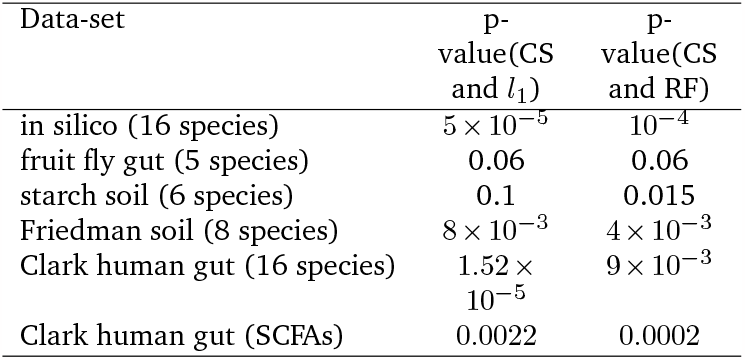
Performance p-values: One-sided permutation test for various data-sets, comparing compressive sensing with a weighted Walsh-Hadamard basis with random forest regression and *l*_1_ -regularized regression.

| Data-set | p-value(CS and $l_1$) | p-value(CS and RF) |
| --- | --- | --- |
| in silico (16 species) | $5 \times 10^{-5}$ | $10^{-4}$ |
| fruit fly gut (5 species) | 0.06 | 0.06 |
| starch soil (6 species) | 0.1 | 0.015 |
| Friedman soil (8 species) | $8 \times 10^{-3}$ | $4 \times 10^{-3}$ |
| Clark human gut (16 species) | $1.52 \times 10^{-5}$ | $9 \times 10^{-3}$ |
| Clark human gut (SCFAs) | 0.0022 | 0.0002 |

We will label each possible sub-community one can obtain from a pool of *S* species using a binary vector 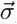, with ones (zeros) denoting the presence (absence) of each species in the initial community. There are 2^*S*^ distinct values of 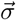 corresponding to the different species combinations possible, with one ecologically trivial case of all species being absent initially. The steady-state abundance of a species *i* in one of the sub-communities, 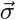, can then be written as 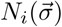, since the abundance by assumption depends on the presence of species in the seed community. Further, since species *i* is absent from half of the 2^*S*^ possible combinations, 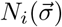 can take up to 2^*S−*1^ non-zero, and potentially distinct, values. Thus, representing species abundances in the 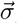 basis requires a binary-ordered vector 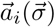, of up to 2^*S−*1^ non-zero coefficients, 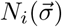. For each species, we have thus defined a map between the 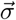 space and the space of steady-state abundances. This formalism, defining maps based on composition, has recently been explored in the context of composition and community functions, where the maps are referred to as ecological landscapes [43– 45]. Here, we define a distinct ecological landscape for each species in the community, where the mapping of interest is between the initial composition of the community, and the final, steady-state abundance of each focal species. Many functions and services of microbial communities depend on species abundances, and thus reliably predicting late-time abundances is a key step on the way to predicting community function.

A promising approach to overcoming this combinatorial challenge would be if there were a more efficient representation of steady-state species abundances using fewer coefficients. In the field of signal processing, we would say that this is possible for a given signal if there is a representation that compresses it. We are not guaranteed that such a representation exists for microbiomes, but below we will show that viewed in the right basis, microbiomes are highly compressible.

## Results

### Species abundances can be represented using only a few coefficients in a novel basis

The question of mapping initial community composition to species abundances at late times is related to the problem of mapping genotypes to fitness in evolutionary genetics, where the effect of interactions between species, which we term ‘landscape interactions’, is analogous to the effect of epistasis among distinct genetic mutations. This connection can made more concrete by considering a combinatorially complete fitness landscape of a genome with *S* positions. The different states of this genome can be enumerated by the presence or absence of a mutation at each of the positions. For each of these 2^*S*^ genotypes, there exists a scalar value of interest, the fitness. The landscape can be represented as a vector 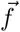, with length, 2^*S*^. Each element is a fitness value, and corresponds to a genotype ordered by an *S*-bit binary number whose digits 1 and 0, respectively, signal the presence or absence of the mutation at the corresponding position [46, 47]. In the case of microbiomes and under the assumptions we outlined in the previous section, the various genotypes are analogous to the distinct sub-communities that are possible when combining species based on initial presence-absence, and the fitness vectors are analogous to the steady-state abundances, with one vector for each species.

Here, we draw upon recent progress in the field of evolutionary genetics to inspire a new approach to microbiome prediction. Specifically, some empirical fitness landscapes are sparse when represented in a type of high-dimensional Fourier basis, called the Walsh-Hadamard basis [47–52]. Moreover, in the field of signal processing, it has been noted that most natural images and many time-series signals are stored efficiently in the same kind of basis [53]. Discovery of these sparse representations, or sparse coding, has then been used to learn entire datasets from sampling only limited data using algorithms from the field of compressive sensing [48, 52, 54, 55].

These breakthroughs in other fields motivated us to consider the representation of species abundances in various sub-communities in a similarly-transformed perspective. Specifically, we used a weighted Walsh-Hadamard basis (Methods and SI) [52, 56, 57]. Up to multiplication by a diagonal matrix, this transform is the well-known Walsh-Hadamard (WH) transform, an analog of Fourier transforms in this discrete basis. It has several useful properties as discussed in the SI, including orthogonality and symmetry [46, 47, 58]. In this work, we denote this weighted Walsh-Hadamard basis as 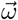. This is different from the presence-absence basis, which we denote using 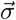. In terms of the abundance of a species in the presence-absence basis, which is given by 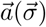, the transformed abundances are given by 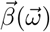 with the following relation between the two representations:

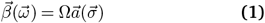

where Ω is the transformation matrix defined as Ω = *V H, H* is the Hadamard matrix and *V* is a diagonal weighting matrix, both described in Methods. Multiplication by the Hadamard matrix *H* transforms the representation into an orthogonal basis, while multiplication by the weighting matrix *V* implements an ecologically motivated assumption that down-weighs higher-order landscape interactions. This is based on the assumption that while higher-order landscape interactions are allowed, lower-order interactions are relatively more likely, and the weighting encoded in *V* captures this expectation. (See Methods and SI Figs. S3 & S7 for more details.)

To test whether microbiomes could plausibly be sparse in the 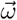 basis, we first generated abundance data from simulated microbial communities using microbial consumer-resource (MiCRM) models [59, 60]. We consider a community of 16 species, with inter-specific interactions mediated by consumption of resources. As elaborated in the Methods section, the MiCRM is a realistic choice for studying in silico microbiomes since its dynamical equations include terms for both cross-feeding and competition. In these in silico communities, species are distinguished by the differences in their ability to consume and secrete various resources. The advantage of starting with simulated communities is that we are able to comprehensively generate all possible species combinations, and thus test rigorously to what extent sparsity does or does not hold in the 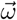, basis. In particular, from a pool of 16 species, we considered the stationary points of the MiCRM corresponding to each of the total possible 2^16^ = 65, 536 species assemblages, obtaining the late-time, steady-state abundances arising from each initial species combination. To ensure the robustness of our conclusions, we considered 10 different sets of 16-species communities.

Figure 2B shows the coefficients in the 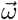, basis, 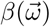, calculated from the steady-state abundances of a typical species from this simulated pool. A small fraction of the 32,768 (= 2^16*−*1^) coefficients are significantly larger than the others and using just the 50 largest coefficients to estimate the observed abundances in each community, we explain 99% of the observed variance (see Figure 2C). Figure 2D shows the goodness of fit as we change the number of 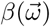 coefficients used to fit species abundance, 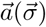. The goodness of fit is measured as a prediction score [52, 61],

**Fig. 1.**
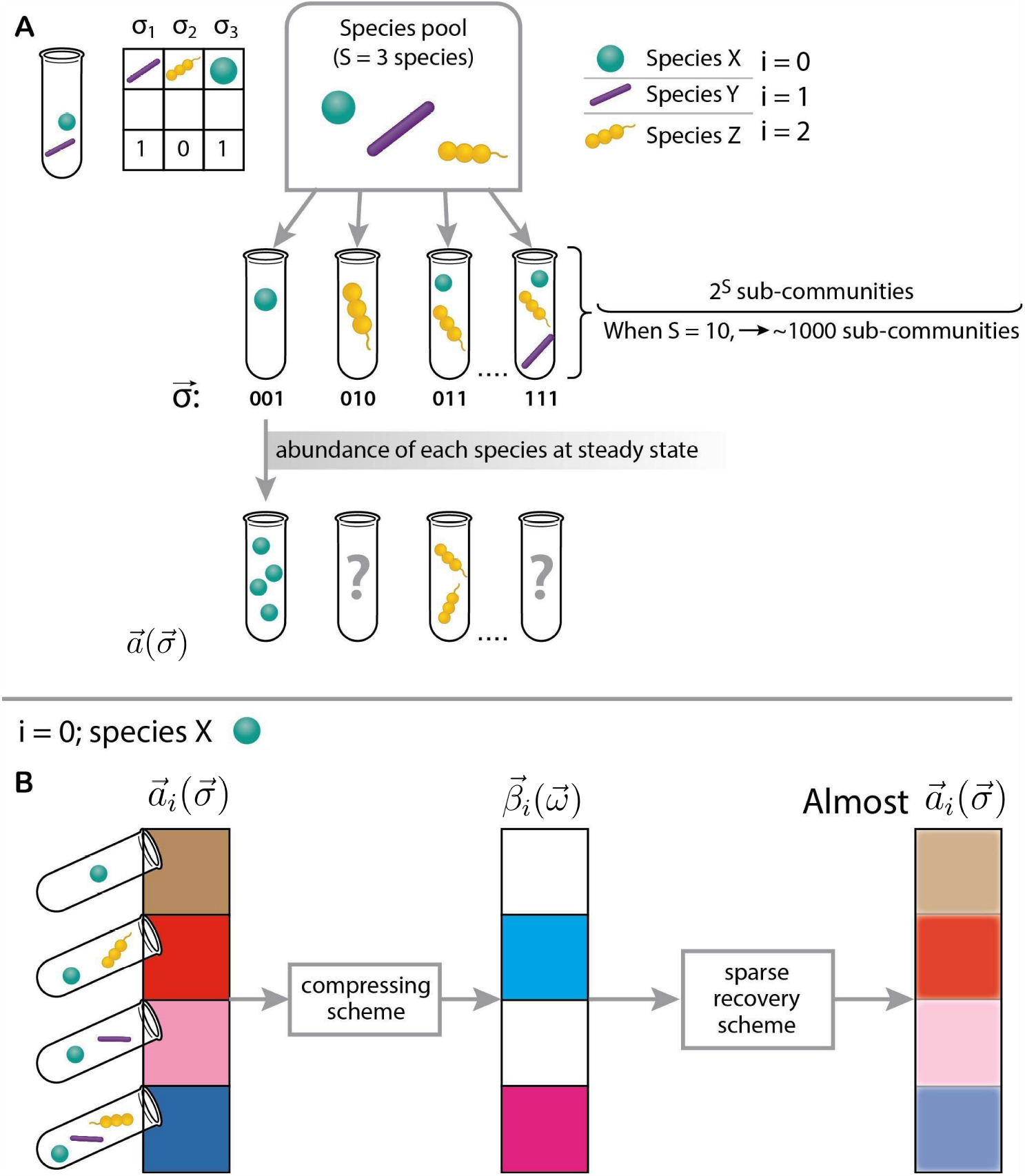
A sparse recovery algorithm can be used to predict microbial community end points. **A** Given a species pool with *S* members, there are 2^*S*^ *−* 1 sub-communities, based on different choices of initial presence-absence of the members. This problem of exponential scale means that only a few sub-communities can be sampled in the laboratory. In this paper, we focus on predicting end-point community compositions of unseen sub-communities, given limited data. **B**. We found that the relative abundances of any species in the pool when stacked according to the sub-communities in which it was initially present is sparse in a weighted Walsh-Hadamard basis 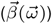. This means that even though a chosen focal species (green, round) may have numerous, different steady-state abundances in different sub-communities, a weighted Walsh-Hadamard representation of this vector will have only a few components. The numerous, different possible steady state abundances are represented by the colors in the column corresponding to 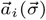. Sparsity of the representation, 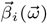, is visualized by the number of transparent components, or boxes, which indicate that a lot of the coefficients in this representation are insignificant. This sparsity may be leveraged by using a sparse recovery technique, called compressive sensing, which prescribes that a much smaller, generic sampling of sub-communities is required to efficiently predict steady-state abundances of the species in all other, unsampled sub-communities.

**Fig. 2.**
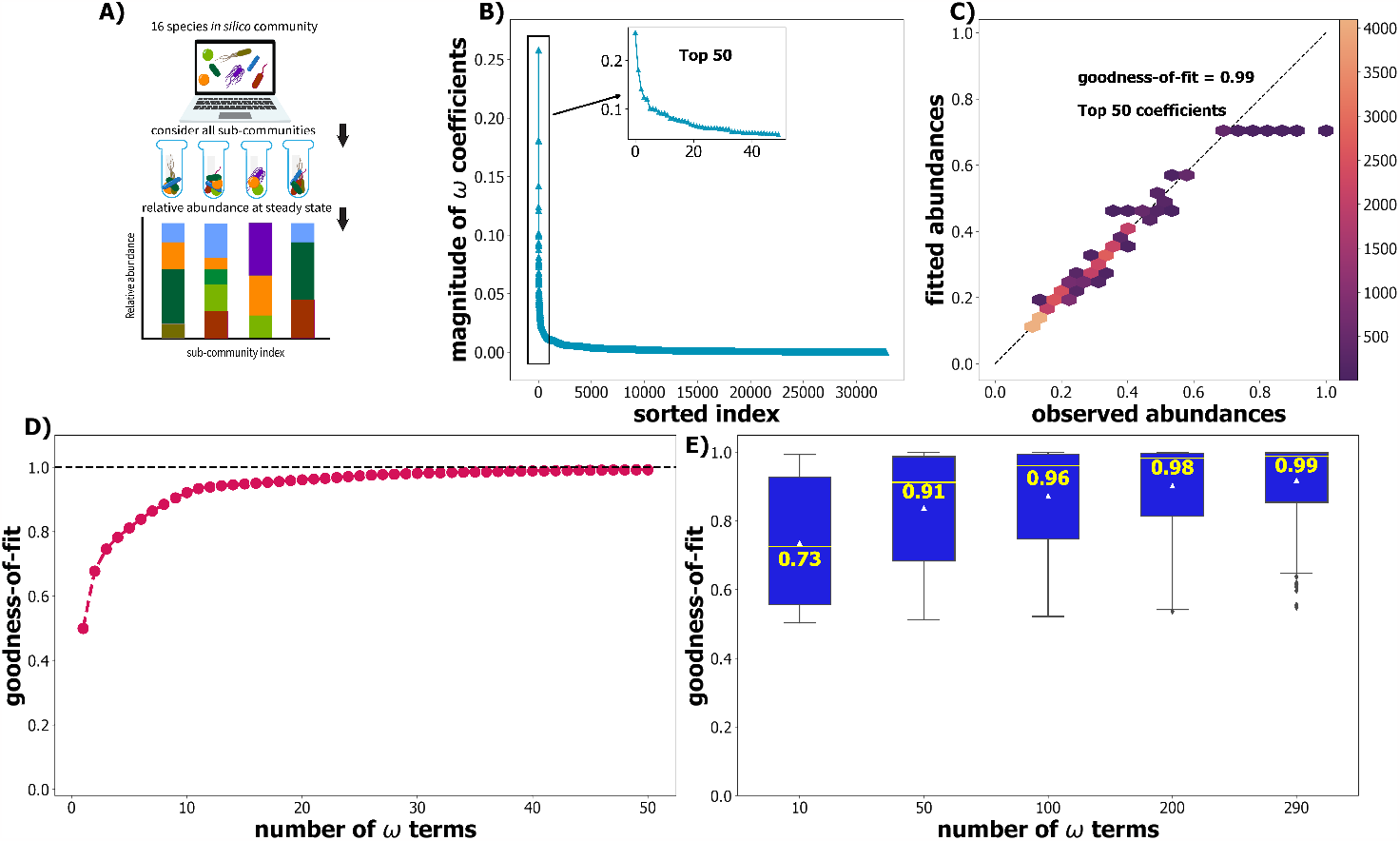
Relative abundances of a simulated community are sparse in the weighted Walsh-Hadamard 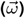 basis. **A)** We simulated a pool of 16-species in silico using microbial consumer-resource models [59, 60]. Simulations started from all 65535 possible sub-communities corresponding to distinct species presence-absences 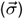 to obtain species abundances in the different steady-state communities. The relative abundances of a species, in the 32678 sub-communities in which it was initially present, constitute the coefficients of 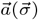. By using a weighted Walsh-Hadamard transform, species abundance can be represented in an orthogonal 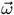 basis. **B)** The coefficients used to represent abundances of a typical species in the 32678 sub-communities using the 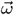 basis are plotted sorted by absolute size. The vast majority of the 32678 coefficients are small; the inset plots the largest 50 coefficients. **C)** Using only 50 coefficients, we were able to explain most (99%) of the variation in species abundances. The color bar indicates the density of points in the hexplot. Panel **D)** demonstrates that a small number of 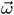 coefficients is sufficient to fit abundances almost perfectly. **E)** Box-plots of goodness-of-fit, aggregated for all 16 species in all 10 replicates, demonstrate that only a small number of 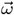 coefficients are required to explain most of the observed variation in abundances. We display medians in the text and by yellow lines, while mean statistics is denoted by white triangles. Coefficients in panel **B)** are ordered by magnitude. The goodness-of-fit is calculated via Equation 2.

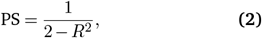

where *R*^2^ is the coefficient of determination. The prediction score always lies between 0 and 1. For a model that could predict only the mean abundance (over all sub-communities in which a species was initially present) regardless of the input sub-community, this prediction score would be 0.5 (corresponding to *R*^2^ = 0). A score of 1 indicates perfect prediction. The high value of this goodness-of-fit measure indicates that we are able to capture most of the variation in species abundance using a very small number of coefficients, reinforcing that this representation of species abundances is highly sparse. This finding is consistent for all species across the 10 species pools we simulated (see Figure 2E).

### Compressive sensing predicts simulated community compositions from limited data

In simulated consumer-resource models, we have established that there is a sparse representation of species abundances. But we obtained this sparse representation from a complete knowledge of all species abundances at steadystate, 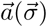, from all initial species combinations. Even if sparsity does also hold in real microbiomes, we will need a method to identify the largest, most significant coefficients of the vector 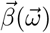 from only limited experimental data. Compressive sensing (CS) provides this method. Assuming that a sparse representation *does* exist for a given signal, compressive sensing will recover the signal in its entirety from only limited, generic sampling [48, 52, 54].

In the context of microbial ecology, results shown in Figure 2 conclusively establish sparsity for our in silico communities: the steady-state abundance vector, 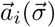, of most species is afforded a sparse rep-resentation in the 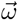, basis. Compressive sensing may then allow us to recover these 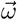, coefficients from lim-ited sampling of abundance data, 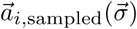. Specifically, using compressive sensing translates to implementing a constrained optimization algorithm that finds a solution, 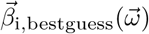, which has the lowest-possible *l*_1_-norm while possessing an inverse Walsh-Hadamard transform that optimally fits the sampled abundances.

In this work, we implemented the Basis Pursuit Denoising (BPDN) algorithm [52, 55, 62] to find the 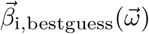. The inverse Walsh-Hadamard transform then yields the model prediction for abundances:

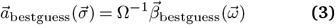

We tested the predictive power of compressive sensing for the in silico communities, for which we have access to all data in practice, and where we have already shown that a sparse basis exists. Specifically, we restricted ourselves to observing/sampling 1% or 327 subcommunities of the 32,678 possible sub-communities with a given species present initially. We also considered lower sampling percentages of 0.1%, 0.2%, and a higher percentage of 2%. With these limited “training” data, we used CS algorithms to estimate coefficients in the 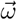, basis. Figure 3A shows that CS is able to infer the most significant coefficients of the ground truth 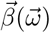 accurately from limited training data, and the accuracy increases as more data are obtained for training.

**Fig. 3.**
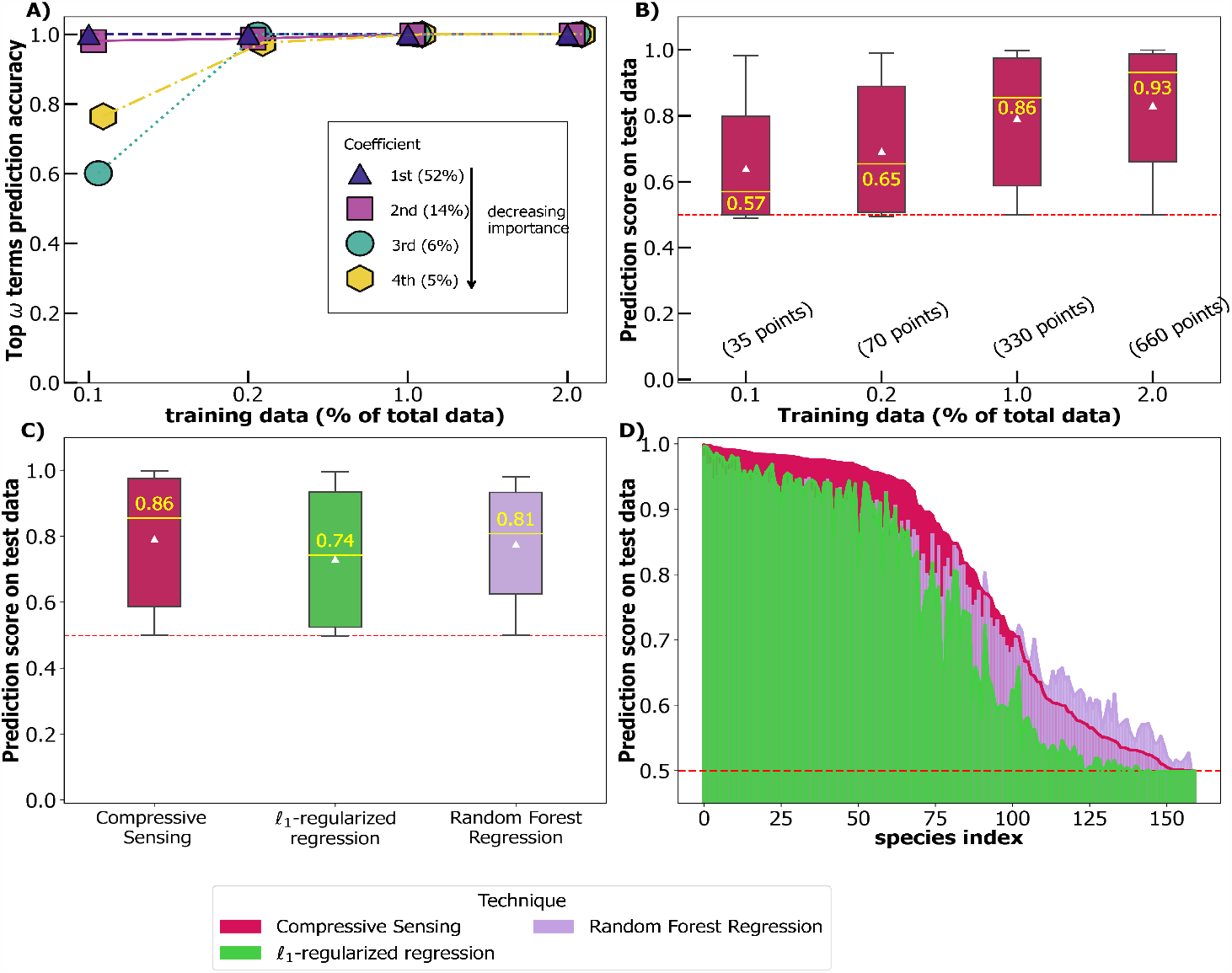
Sparsity in the 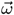 coefficients implies abundances can be practically recovered from limited data in simulated communities. **A)** The accuracy of predicting the 4 most significant *ω* coefficients by compressive sensing (CS) increases as the size of training data is increased. The largest coefficients, which explain most of the variance in abundance, are more easily predicted than smaller coefficients from limited data. The percentages in the parenthesis next to the coefficient symbol and index denote the percentage of the variance explained by each coefficient taken individually. This means, for example, that the first *ω* term explains 52% of the total variance. **B)** The ability of compressive sensing to predict species abundances in new sub-communities when trained on a small fraction of all possible sub-communities is shown using the Prediction Score (equation 2). Note that 1% corresponds to 327 sub-communities, which would requires 4 96-well plates. The median score, shown by the yellow line is annotated. The number of sub-communities sampled at each training level is indicated within parentheses. **C)** The prediction score of two alternative methods, *l*_1_ -regularized regression and random forest regression is shown for comparison. We controlled for data so that all the methods were trained on the same 327 (1%) communities. Box plots denote the performance across the 160 species (10 replicates of pools with 16 species). A model that outputs only the mean abundances of a species in all sub-communities in which the species was initially present, regardless of the input sub-communities, would get a prediction score of 0.5. We see that compressive sensing does objectively better than such a model (prediction score shown using red dashed line) while outperforming random forest regression and *l*_1_ -regularized regression. **D)** Comparing the performance of CS, *l*_1_ -regularised regression, and random forest on a species-by-species basis, we find that some species are easier to predict than others. Nevertheless, CS outperforms both regressions for most species. Permutation test p-value for the difference in the performance at 1% sampling between compressive sensing and random forest regression is statistically significant at 10^*−*4^ and that between compressive sensing and *l*_1_ -regularized regression is 5 *×* 10^*−*5^. All results are averages over 5 runs.

Next, we compare the recovered abundances with the ground-truth abundances. We compute the recovered abundance for a species using Equation 3, where 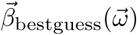 is the Walsh-Hadamard vector inferred by the BPDN algorithm when trained on the limited abundance data for the species. Figure 3B shows the resulting prediction score on out-of-sample abundance data.

We see that the algorithm’s predictive power improves with more training data, and appears to asymptote at as little as 1% of all possible data included in the training set. We compare our method with two alternative approaches: an *l*_1_-regularized regression on the 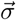 basis that tries to estimate 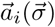 directly, and random forest regression. (See Methods for detailed description of these algorithms) The *l*_1_-regularized regression is adopted since the number of observations is smaller than the number of possible coefficients, i.e., the problem is under-determined. We found that compressive sensing outperforms both regularized regression and random forest regression, as can be seen from the median values of the predictions in Figure 3C. Finally, Figure 3D compares the prediction score of the three methods for each species, which is a matched comparison at the species level of the three approaches, with training data held at 1%. This analysis reveals that some species are easier to predict, with a prediction score close to 1, while some others that are harder to pre-dict. In the SI, we show that species which are easier to predict are dominated by low-order landscape interactions, and have a smoother abundance landscape (see Fig. S4 and ref. [58]). We can thus conclude that prevalence of low-order interactions lead to a more predictable landscape. Despite this variation in performance at the species-level, compressive sensing outperforms *l*_1_-regularized regression in 97% of the cases, and outperforms random forest regression in a majority of the cases (67%).

Thus, compressive sensing provides a way to use the sparse basis for our simulated consumer-resource communities, along with limited observations of species abundances in different sub-communities, to predict species abundances out-of-sample.

### Compressive sensing predicts species abundances in experimental data

Using simulated communities, we demonstrate that the challenge posed by an exponential number of sub-communities to the problem of ecological predictions is massively reduced with compressive sensing. We now test the performance of CS as a predictive tool in a diverse range of real microbiomes, in the context of unknown community dynamics which likely depart from our idealized consumer-resource model, in addition to the difficulties posed by both stochasticity and incomplete data. We draw data from four published studies of microbial communities spanning *in vitro* and *in vivo* conditions, pool sizes from 5 to 16, and microbes from environments including the human gut, soil, and fruit flies [28, 29, 63, 64].

In the first study, Gould *et al*. assembled all combinations of 5 species *in vivo* in the gut of *Drosophila melanogaster*. The remaining three studies considered larger species pool sizes and hence assembled only a fraction of all possible species combinations *in vitro*. Sanchez-Gorostiaga *et al*. assembled 53 combinations of 6 starch-degrading soil microbes. Friedman *et al*. assembled 101 combinations of 8 soil microbes. From Clark *et al*., we consider 187 combinations assembled of 16 gut microbes. Figure 4 demonstrates the performance of compressive sensing, *l*_1_-regularized regression, and random forests on predicting unseen communities using a k-fold cross-validation procedure (see Methods). We used a k-fold cross-validation procedure due to the small number of data points and report the prediction score, at the optimal value of the hyperparamter, on the stacked validation sets. Further, keeping the bias-variance trade-off [65] in mind, we set *k* = 3 for most experimental communities. A complete description is given in Methods. Figure 4 B-E show the performance of compressive sensing, alongside the alternative methods, on the individual species. As in simulations, we find that some species are easier to predict than others. Compressive sensing outperforms regularized regression in the majority of the cases (32 out of 35), and also does better than random forest regression in 29 of 35 cases. At the community level, as shown in Fig. 4, compressive sensing outperforms both methods when comparing the mean prediction score on a data-set. This difference in prediction is statistically significant for the two largest datasets. This is tabulated in Table 1, where we list the p-values of a one-sided permutation test. We also note that species which are harder to predict using compressive sensing remain hard to predict when using the other methods. In summary, our compressive sensing approach enables more accurate predictions of microbial community abundances.

**Fig. 4.**
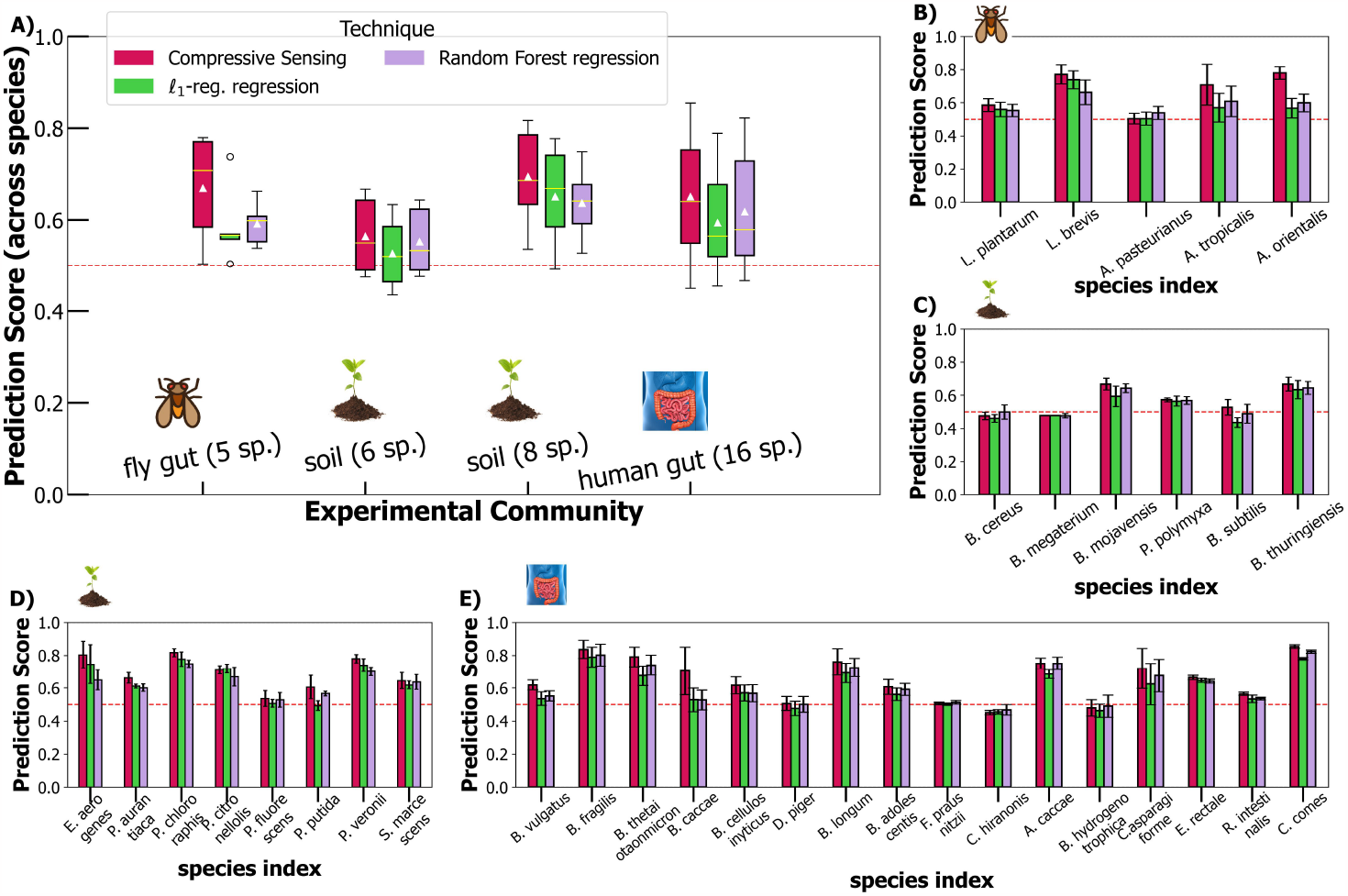
Performance of compressive sensing on 4 experimental data. We tested the performance of compressive sensing in 4 real microbiomes, ranging from 5-16 species, and in both *in vitro* and *in vivo* contexts (Methods). **A)** At a community level, compressive sensing outperforms both random forest regression and *l*_1_ -regularized regression. **B)** For the *in vivo* gut fly dataset from [28, 66], compressive sensing does better in predicting a majority (4 of 5) of the species. Permutation test p-values are .09 and .06 for comparison against random forest regression and *l*_1_ -regularized regression. **C)** For the *in vitro* 6-species soil community [29, 67], we find that compressive sensing outperforms random forest marginally (permutation test p-value : 0.1) but does better than *l*_1_ -regularized regression (p-value : 0.015). This dataset, overall, is harder to predict. **D)** Compressive sensing outperforms both random forest regression and *l*_1_ -regularized regression (pvalues : 4 *×* 10^*−*3^ and 8 *×* 10^*−*3^) in the 8-species *in vitro* soil community assembled by Friedman et al. [63]. At the species level, compressive sensing improves predictions in 7 of 8 cases when compared to *l*_1_ -regularized regression, and in all cases when compared to random forests. **E)**. Using a 16 species community assembled by Clark et al. [64, 68], we find that compressive sensing outperforms other methods reaching statistically significant values in a permutation test (p-value : 9 *×* 10^*−*3^ for comparison with random forest regression, and 1.52 *×* 10^*−*5^ for *l*_1_ -regularized regression). At the species level, compressive sensing improves predictions in 15 of 16 cases when compared to *l*_1_ -regularized regression, and in 12 cases when compared to random forests. We report all results as averages of 5 runs. The error bars in panels **B), C), D)**, and **E)** indicate standard deviation from the mean across 5 runs. Species names on the x-axis are compiled from the data as provided by authors of the studies.

**Fig. 5.**
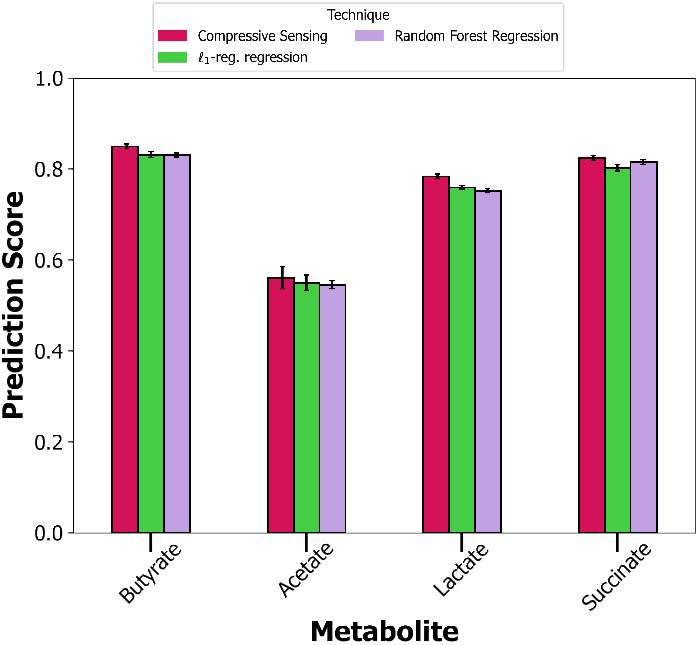
Compressive sensing is able to directly predict community functions. Along with abundances of species, Clark et al [64, 68], reported concentrations of 4 organic acid fermentation products: butyrate, acetate, lactate, and succinate. Of these, butyrate is reported to be particularly important to human health. For these important community functions of metabolite production, we find that compressive sensing does well with *PS >* 0.5, and also outperforms the methods of *l*_1_ -regularized regression and random forest regression. Permutation p-values for difference in means between compressive sensing and *l*_1_ -regularized regression and that between compressive sensing and random forest regression indicate statistical significance (p-values : 0.0022 and 0.0002). We report averages over 3 runs. Cross-validation *k* fold was chosen to be 10.

## Discussion

A central goal of ecology is to understand emergent properties of microbiomes. In particular, we want to be able to understand and quantitatively predict outcomes of community assembly. Furthermore, community composition may be predictive of community function, and systematically optimizing microbial community function is a critical goal in microbiome engineering. However, an exhaustive search through all possible ways to combine microbial taxa is impractical, and fitting mechanistic models can still require large amounts of data, while also imposing assumptions about community dynamics that may fail to hold for real microbiomes. Here, we establish the method of compressive sensing as a new, model-agnostic, predictive tool, applicable in situations where experimenters have access to limited data. The success of our approach relies on the assumption that species abundances are sparse in a novel basis, which we demonstrate explicitly for simulated consumer-resource models, and test in experimental data. While this kind of sparsity appears in many areas of signal processing, and has also been applied in an evolutionary genetics context [52–54, 69], the different and complex dynamics underlying microbiome communities means that this sparsity did not *a priori* need to hold in microbiomes. The fact that it does may open up new avenues for assembling microbial communities and optimizing their function.

We use this method *in silico, in vivo*, and *in vitro* to predict final steady-state abundances of all species. We also show that our method outperforms *l*_1_-regularization and a widely-used machine learning method, random forest regression. We also find that the sparsity of microbial landscapes is not limited to their late-time, steady-state abundances, but can be extended to community functions. In Fig.5, we show that this method can also robustly predict community functions: for the 16-species data-set studied in [64], compressive sensing can be used to accurately predict the amount of butyrate, succinate, lactate, and acetate produced by combinations of these species. By leveraging the interpretability of Walsh-Hadamard coefficients and the roughness of the community-function landscape, we also found that butyrate production, in particular, could be associated with two key species, *Desulfovibrio piger* (DP) and *Anaerostipes caccae* (AC). The absence of DP was found to increase butyrate production, while the absence of AC decreased it. This inference matches what has been found in the study in [64]: hydrogen sulfide production by DP inhibits butyrate production. This reinforces a key assumption, that emergent properties of microbiomes, perhaps even incredibly complex community-level functions, may be thought of as effectively arising from only a few degrees of freedom. On the side of microbiome engineering, this result is useful in practice — using sparsely sampled structure-function landscapes, we may be able to recover entire landscapes and hence reliably look for optimal functions and corresponding communities without taking into account intricate mechanistic models. This formalism does not require time-series data, making it easier to work with data obtained from high-throughput laboratory techniques. Further, it performs well with relative abundances, which is in line with how abundances are widely observed and calculated in this field, though in principle the approach will also work well on absolute abundances (See SI Figs. S8 and S9).

Recently, there has been work on applying deep-learning models to predict community structures [32, 34], providing a natural point of comparison with our approach. These models can predict well, but their performance can be obscured by large numbers of hyper-parameters and a difficulty in interpretability. On the contrary, our method is naturally endowed with interpretation in terms of the sparseness of higher-order landscape interactions, and the compressive sensing algorithm requires only one tunable hyper-parameter. In cases where compressive sensing predicts species abundances accurately, it is reasonable to conclude that higher-order interactions are sparse, and that low-order interactions dominate. Our approach therefore provides a kind of middle-ground between mechanistic models, which may be challenging to parametrize well, and purely statistical models, which are hard to interpret. The identification of sparsity renders the nature of the landscape in real communities both more predictable and better interpretable.

While our approach works well in the simulated and experimental data we applied it to, this method hinges on the assumption that there is a unique steady-state set of community abundances, rather than the potential for multiple equilibria, or more complex late-time dynamics [35–38, 63]. There is therefore scope to consider further generalizations in cases where experimental data suggests that these outcomes are possible. However, for cases where experiment suggests a unique map from initial composition to late-time abundances, our approach paves a new way to reliably engineer microbial communities.

## Methods

### Representation of Abundances in the Wal-sh-Hadamard Basis

We consider a combinatorially complete data-set of microbial abundances at steady state. Starting with a set of *S* species, there are 2^*S*^ − 1 of combining them, each assemblage characterised by the species initially present. We consider that the underlying population dynamics of these sub-communities, whatever they are, give rise to a unique steady state for each sub-community, i.e we do not consider cases of multi-stability. Borrowing from the language of genetics and epistasis, we consider a vector of abundances at steady state for each of the *S* species. For concreteness, in this section, we consider a set of 3 species. The set of all the sub-communities, each with a different starting composition is enumerated as: 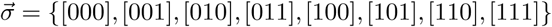. We include the ecologically trivial case of all species being initially absent. The element [011], for example, denotes the sub-community with species 2 and 3 present at the start. For each of these unique conditions, we consider the steady-state abundance (say, cell counts) of each species. This allows for a mapping to be written down explicitly, between the decimal-ordered binary elements and the *S* steady-state abundance vectors.

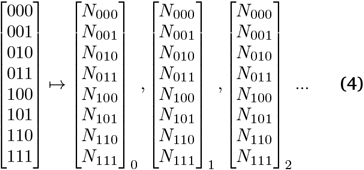

Here the subscripts on the vectors denote the species indices. For each species, we can further build 2^*S−*1^-long vectors, since every species is initially present in only the half the total possible 2^*S*^ sub-communities. We can define a Walsh-Hadamard transform on this vector of steady-state abundances of a species *i*. In particular, we work with a weighted Walsh-Hadamard transform. The weighted transform is implemented by the matrix: *V H* where the matrices *V* and *H* are generated by the recursion relations:

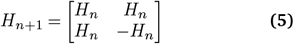

with *H*_0_ = 1

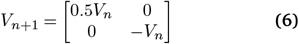

with *V*_0_ = 1.

*V* is a diagonal weighting matrix that takes into account the order of interactions, to account for averaging over different numbers of terms as a function of the order of interactions [56]. In our case, it serves to provide a way to bias inference from limited data towards lower-order interactions. (See SI Section 5 and Fig. S7)

### Compressive Sensing and Sparse Recovery Algorithms

Implementation of compressive sensing involves using optimization algorithms that find a sparse representation of the data using small subsets of observations. After a sparse representation has been found by the algorithm, the rest of the unseen data is computed by taking the inverse transform of the best-guess sparse representation. This inverse transform is the inverse of the matrix, Ω, that is expected to sparsify the data. To infer the sparse representations from limited data, we employed the basis pursuit denoising (BPDN) algorithm [55] which is an optimization problem posed as: 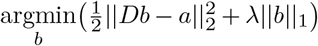. This is the same as LASSO (Least Absolute Shrinkage and Selection Operator) [53]. The idea is to find the *b* vector that has the smallest *l*_1_-norm while keeping the error from observed data as low as possible. To find the sparsest representation, the problem should actually minimize the *l*_0_-norm instead of *l*_1_, but this problem is non-convex and combinatorial [53], and *l*_1_-norm minimization is a convex approximation to this. This algorithm has only one hyperparameter, *λ*, which can be tuned according to the expected sparsity of the representations. In real datasets, this value needs to be chosen carefully. We report all results in the main text based on the optimal hyperparameter. We used the SPORCO package [70] in Python to implement BPDN using an alternating direction method of multipliers (ADMM) algorithm [71].

#### Problem set-up

Given a S-species combinatorially incomplete data-set with *n* different sub-communities sampled, we want to predict the abundances of species outside out of the sampled experiments. We focus on a species, *i*, for which we have *n* sampled abundances, and we make predictions for the remaining 2^*S−*1^ − s*n* sub-communities. We assume, for most species in the pool, that the abundances, 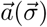, are sparse in the 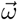, basis, i.e 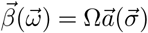 is a sparse vector, with only a few significant coefficients.

#### Algorithm implementation

We implement the BPDN/LASSO algorithm with 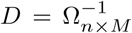, here *D* is the partial Ω^*−*1^ matrix with rows chosen such that the row indices correspond to the decimal representation of the sampled *n* sub-communities, and *M* = 2^*S−*1^. The algorithm then returns the optimal 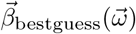 which is a solution to the program:

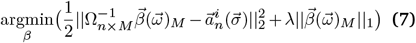

Using Equation 3 of the main text, we recover the complete 2^*S−*1^-long abundance vector for species *i*. For the in silico data-set, we compute the prediction score on the out-of-sample data, whose ground truth is known, using Equation 2:

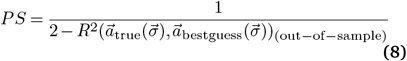

This procedure is repeated for all species in the pool, each with its own set of sampled sub-communities. For experimental datasets, with only partial abundance vectors available, we use *k*−fold CV, by dividing the available data-set for each species into *k*−folds, and solving the program in Equation 7 for sampled abundances in the training folds, and computing the prediction score on the stacked abundances corresponding to the data-points in the validation folds.

#### *l*_1_**-regularized regression**

For a species *i* with relative abundances 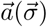, we consider a representation

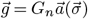

Here *G* is a matrix defined recursively as :

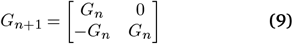

with *G*_0_ = 1 We note that, as elucidated in [52, 56], this transformation can be thought of as looking at the abundance landscape of a species as a local approximation around a single sub-community — the one with all species absent. This is akin to a Taylor expansion on the landscape, while the class of Walsh-Hadamard transforms corresponds to a Fourier transform [72] on the landscape, and is an approximation over the background of all sub-communities. We implement a similar BPDN algorithm as for compressive sensing with Walsh-Hadamard transform, where the algorithm tries to learn the *g* coefficients.

#### Bench-marking with Random Forest Regression

Tree-based ensemble learners, like the random forest regressor and xgboost, are popular choices of supervised learning algorithms, especially when the predictors are not sure to be linearly related to the target variables. We implemented the random forest regressor as available in scikit-learn in Python [73]. While there are arguments for using a multi-output regression instead of several single-output ones [74], we worked with fitting a random forest model to each species individually. For data control and an apples-to-apples comparison, we used the same data splits and number of folds as we did in compressive sensing. With random forest regression, there are multiple hyper-parameters to consider. To reduce the number of hyper-parameters to tune, given the small size of some of the experimental datasets, we used the number of estimators (trees) to be the scikit-learn default (100) and the number of features for selection at every split to be 1*/*3 of the number of species in each case.

#### Predictions on a 5-species gut community

Gould *et al*. [28] assembled all possible communities from a 5-species pool of bacteria that are known to colonize fly guts in both wild and laboratory conditions. All the 32 sub-communities (termed treatments) were assembled in germ-free flies, each with 48 replicate experiments. They reported colony-forming unit (CFU) counts for each replicate and each treatment. For each replicate, we first computed the relative abundance of each species in that community and then took the average of these relative abundances of each condition (i.e each sub-community type). Complete data for this study is available in Ref. [66].

#### Predictions on a 6-species starch-degrading community

Sanchez-Gorostiaga *et al*.[29] studied assemblages by considering a pool of 6 amylolytic soil bacterial species. Out of the 63 possible communities, they reported abundance data (in colony-forming units) for 53 communities. We dropped two communities that were inconsistent with labeling, and worked with 51 data points. For each community and each replicate, we calculated the total biomass of the community, and divided the abundance of each species to find the relative abundance of each species in the community. If multiple replicate experiments were reported, we first computed the relative abundance of a species in each replicate and then averaged this across replicates to find the mean relative abundance. Data for this study is available in Ref. [67].

#### Predictions on an 8-species soil microbiome community

Friedman *et al*. [63] studied a community of 8 heterotrophic soil-dwelling bacteria, reporting their optical densities (ODs) after cross-checking actual cell counts using agar plating. They considered sub-communities which included all the 8 monocultures, all pairwise combinations, all three-species communities, all 8 leave-one-out communities, and the sub-community with all the species present. They combined species in different ratios (i.e other than 1 : 1), and sampled the populations at various times, obtaining time-series data. This difference in starting abundances however did not affect the steady-state in most conditions and replicates. However, for 2 sub-communities they found that the steady-state abundances within the replicates had a much larger variation than what could be accounted for by experimental noise, and one sub-community displayed bistability. We discarded these experimental conditions, and for all other data points proceeded to take the average of the final time (where we assume steady state has been attained) abundances across the replicates. Data for this study is available in Ref. [75].

#### Predictions on a subset of a 25-species synthetic human gut community

Clark *et al*. [64] studied a community of 25 species that consisted of species spanning all major phyla in the human gut microbiome, and also are representative of the major metabolic functions in the gut. Of the 2^25^ − 1 ecological communities possible, they sampled ∼ 600. For these sub-communities, they reported read counts, and computed the relative abundances for each species using total read counts for each sub-community. They also found absolute abundances by multiplying the relative abundance with the OD_600_ measurement for each sample. As outlined in the Methods section of [64], we excluded sub-communities that were flagged as contaminated. Data for this study is available in Ref. [68]. In such a data-limited case, the performance of any algorithm will be difficult to test. Therefore, we considered a sub-set of 16 species with the other 9 always being absent. Since there are many possible sub-sets of 16 species in the background of any consistently-absent 9 species, we selected a 16-species pool such that the number of experimental data-points available to us was maximized. The distribution of available data-points with different number of species consistently absent is shown in SI Fig. S10. With this, we had ∼ 0.2% of this 16-species landscape available to us. We considered relative abundances in the data, after averaging over the number of replicate experiments for each sub-community. There were differing number of data points corresponding to each species, unlike the previous data-sets. We therefore allowed *k* to vary in the cross-validation scheme. *k* was set to 3, 5, or 7 depending on the size of the data-set for each species. The authors also reported concentrations of 4 organic acid fermentation products: butyrate, acetate, lactate, and succinate. For these community functions of metabolite production, we used a 10-fold cross-validation approach.

#### Microbial consumer-resource models

We considered a pool of 16 species, with a single externally supplied resource in a chemostat. Communities with cross-feeding that are supplied with a single resource externally have been studied in recent experiments [76–80]. In our simulations, the supplied resource is primarily consumed by only a few species. However, in the presence of crossfeeding, there are 20 metabolites present in the community. Different species consume different resources at different rates, this information is encoded in the consumer matrix, *C*, with elements *c*_*iα*_, which characterizes a kind of pairwise interaction, but here between consumer and resource, rather than directly between pairs of species. Such pairwise consumer-resource interactions can give rise to both pairwise interspecific interactions and higher-order interspecific interactions [81]. We choose a consumer matrix whose elements are sampled from a binary-gamma distribution [44]. We chose a binary-gamma distribution because this allows for positive uptake rates, but also allows us to control the sparsity of the consumer matrix. By considering all possible combinations of initial presence-absence of the species, we generated steady state abundances of all the species, by numerically solving the ODEs for the consumer-resource model (equations given in SI section 3). Simulations reached steady-state when the root mean square of the logarithmic growth rates rates of the species fell below a threshold, 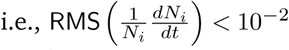. Further, we verified that the extinct species could not have survived in the community by simulating a re-invasion attempt of the steady-state community.

Thus, for each species, we have 2^16^ abundance data points. Since each species is present in only half the combinations, the effective total data for each species is 2^15^. Further, we considered the relative abundance of a species in each combination. We calculated the relative abundance by summing the biomass of each species in the sub-community and dividing this by the total biomass of that sub-community. Details of parameters used in the simulations are given in Section 3 of the SI.

## Bibliography

1. Falkowski, P. G., Fenchel, T. – Delong, E. F. The Microbial Engines That Drive Earth’s Biogeochemical Cycles. Science 320, 1034–1039 (2008). URL https://www.sciencemag.org/lookup/doi/10.1126/science.1153213.

2. Paerl, H. W. – Pinckney, J. L. A Mini-Review of Microbial Consortia: Their Roles in Aquatic Production and Biogeochemical Cycling. Microbial Ecology 31, 225–247 (1996). URL https://www.jstor.org/stable/25152978.

3. Flint, H. J., Scott, K. P., Duncan, S. H., Louis, P. – Forano, E. Microbial degradation of complex carbohydrates in the gut. Gut Microbes 3, 289–306 (2012). URL http://www.tandfonline.com/doi/abs/10.4161/gmic.19897.

4. Brenner, K., You, L. – Arnold, F. H. Engineering microbial consortia: a new frontier in synthetic biology. Trends in Biotechnology 26, 483–489 (2008). URL https://linkinghub.elsevier.com/retrieve/pii/S0167779908001716.

5. Olle, B. Medicines from microbiota. Nature Biotechnology 31, 309–315 (2013). URL http://www.nature.com/articles/nbt.2548.

6. Tanoue, T. et al. A defined commensal consortium elicits CD8 T cells and anti-cancer immunity. Nature 565, 600–605 (2019). URL http://www.nature.com/articles/s41586-019-0878-z.

7. Minty, J. J. et al. Design and characterization of synthetic fungal-bacterial consortia for direct production of isobutanol from cellulosic biomass. Proceedings of the National Academy of Sciences 110, 14592–14597 (2013). URL http://www.pnas.org/lookup/doi/10.1073/pnas.1218447110.

8. Lindemann, S. R. et al. Engineering microbial consortia for controllable outputs. The ISME Journal 10, 2077–2084 (2016). URL http://www.nature.com/articles/ismej201626.

9. Buffie, C. G. et al. Precision microbiome reconstitution restores bile acid mediated resistance to Clostridium difficile. Nature 517, 205–208 (2015). URL http://www.nature.com/articles/nature13828.

10. Hu, J. et al. Design and composition of synthetic fungal-bacterial microbial consortia that improve lignocellulolytic enzyme activity. Bioresource Technology 227, 247–255 (2017). URL https://linkinghub.elsevier.com/retrieve/pii/S096085241631728X.

11. Senne de Oliveira Lino, F., Bajic, D., Vila, J. C. C., Sánchez, A. – Sommer, M. O. A. Complex yeast–bacteria interactions affect the yield of industrial ethanol fermentation. Nature Communications 12, 1498 (2021). URL https://www.nature.com/articles/s41467-021-21844-7.

12. Weng, J.-K., Li, X., Bonawitz, N. D. – Chapple, C. Emerging strategies of lignin engineering and degradation for cellulosic biofuel production. Current Opinion in Biotechnology 19, 166–172 (2008). URL https://www.sciencedirect.com/science/article/pii/S0958166908000268.

13. Piccardi, P., Vessman, B. – Mitri, S. Toxicity drives facilitation between 4 bacterial species. Proceedings of the National Academy of Sciences 116, 15979–15984 (2019). URL https://www.pnas.org/doi/abs/10.1073/pnas.1906172116.

14. Swenson, W., Wilson, D. S. – Elias, R. Artificial ecosystem selection. Proceedings of the National Academy of Sciences 97, 9110–9114 (2000). URL http://www.pnas.org/cgi/doi/10.1073/pnas.150237597.

15. George, A. B. – Korolev, K. S. Ecological landscapes guide the assembly of optimal microbial communities. PLOS Computational Biology 19, e1010570 (2023). URL https://journals.plos.org/ploscompbiol/article?id=10.1371/journal.pcbi.1010570.

16. May, R. M. Will a large complex system be stable? Nature 238, 413–414 (1972).

17. Maynard, D. S., Miller, Z. R. – Allesina, S. Predicting coexistence in experimental ecological communities. Nature Ecology – Evolution 4, 91–100 (2020). URL http://www.nature.com/articles/s41559-019-1059-z.

18. Skwara, A., Lemos-Costa, P., Miller, Z. R. – Allesina, S. Modelling ecological communities when composition is manipulated experimentally. Methods in Ecology and Evolution 14, 696–707 (2023). URL https://onlinelibrary.wiley.com/doi/abs/10.1111/2041-210X.14028.

19. Barbier, M., Arnoldi, J.-F., Bunin, G. – Loreau, M. Generic assembly patterns in complex ecological communities. Proceedings of the National Academy of Sciences 115, 2156–2161 (2018). URL https://www.pnas.org/doi/full/10.1073/pnas.1710352115. Publisher: Proceedings of the National Academy of Sciences.

20. Allesina, S. – Tang, S. Stability criteria for complex ecosystems. Nature 483, 205–208 (2012). URL https://www.nature.com/articles/nature10832. Number: 7388 Publisher: Nature Publishing Group.

21. Bunin, G. Ecological communities with Lotka-Volterra dynamics. Phys. Rev. E 95, 042414 (2017). URL http://link.aps.org/doi/10.1103/PhysRevE.95.042414.

22. Grilli, J. et al. Feasibility and coexistence of large ecological communities. Nat Commun 8, 14389 (2017). URL https://www.nature.com/articles/ncomms14389. Number: 1 Publisher: Nature Publishing Group.

23. Barabás, G., J. Michalska-Smith, M. – Allesina, S. The Effect of Intra- and Interspecific Competition on Coexistence in Multispecies Communities. The American Naturalist 188, E1–E12 (2016). URL https://www.journals.uchicago.edu/doi/full/10.1086/686901. Publisher: The University of Chicago Press.

24. Bunin, G. Interaction patterns and diversity in assembled ecological communities (2016). URL http://arxiv.org/abs/1607.04734. ArXiv:1607.04734 [cond-mat, physics:physics, q-bio].

25. Serván, C. A., Capitán, J. A., Grilli, J., Morrison, K. E. – Allesina, S. Coexistence of many species in random ecosystems. Nat Ecol Evol 2, 1237–1242 (2018). URL https://www.nature.com/articles/s41559-018-0603-6. Number: 8 Publisher: Nature Publishing Group.

26. Serván, C. A. – Allesina, S. Tractable models of ecological assembly. Ecology Letters 24, 1029–1037 (2021). URL https://onlinelibrary.wiley.com/doi/abs/10.1111/ele.137020. _eprint: https://onlinelibrary.wiley.com/doi/pdf/10.1111/ele.13702.

27. Ansari, A. F., Reddy, Y. B. S., Raut, J. – Dixit, N. M. An efficient and scalable top-down method for predicting structures of microbial communities. Nature Computational Science 1, 619–628 (2021). URL https://www.nature.com/articles/s43588-021-00131-x.

28. Gould, A. L. et al. Microbiome interactions shape host fitness. Proceedings of the National Academy of Sciences 115, E11951–E11960 (2018). URL http://www.pnas.org/lookup/doi/10.1073/pnas.1809349115.

29. Sanchez-Gorostiaga, A., Bajić, D., Osborne, M. L., Poyatos, J. F. – Sanchez, A. High-order interactions distort the functional landscape of microbial consortia. PLOS Biology 17, e3000550 (2019). URL https://dx.plos.org/10.1371/journal.pbio.3000550.

30. Bairey, E., Kelsic, E. D. – Kishony, R. High-order species interactions shape ecosystem diversity. Nature Communications 7, 12285 (2016). URL https://www.nature.com/articles/ncomms12285.

31. Morin, M. A., Morrison, A. J., Harms, M. J. – Dutton, R. J. Higher-order interactions shape microbial interactions as microbial community complexity increases. Scientific Reports 12, 22640 (2022). URL https://www.nature.com/articles/s41598-022-25303-1.

32. Baranwal, M. et al. Recurrent neural networks enable design of multifunctional synthetic human gut microbiome dynamics. eLife 11, e73870 (2022). URL https://elifesciences.org/articles/73870.

33. Fisher, C. K. – Mehta, P. Identifying Keystone Species in the Human Gut Microbiome from Metagenomic Timeseries Using Sparse Linear Regression. PLoS ONE 9, e102451 (2014). URL https://dx.plos.org/10.1371/journal.pone.0102451.

34. Michel-Mata, S., Wang, X.-W., Liu, Y.-Y. – Angulo, M. T. Predicting microbiome compositions from species assemblages through deep learning. iMeta 1, e3 (2022). URL https://onlinelibrary.wiley.com/doi/abs/10.1002/imt2.3.

35. Amchin, D. B., Martínez-Calvo, A. – Datta, S. S. Microbial mutualism generates multistable and oscillatory growth dynamics. bioRxiv 2022.04.19.488807 (2022). URL https://www.biorxiv.org/content/10.1101/2022.04.19.488807v1.

36. Amor, D. R., Ratzke, C. – Gore, J. Transient invaders can induce shifts between alternative stable states of microbial communities. Science Advances 6, eaay8676 (2020). URL https://www.science.org/doi/10.1126/sciadv.aay8676.

37. Debray, R. et al. Priority effects in microbiome assembly. Nature Reviews Microbiology 20, 109–121 (2022). URL https://www.nature.com/articles/s41579-021-00604-w.

38. Björk, J. R. et al. Synchrony and idiosyncrasy in the gut microbiome of wild baboons. Nature Ecology – Evolution 6, 955–964 (2022). URL https://www.nature.com/articles/s41559-022-01773-4.

39. Hastings, A., Hom, C. L., Ellner, S., Turchin, P. – Godfray, H. C. J. Chaos in ecology: is mother nature a strange attractor? Annual review of ecology and systematics 24, 1–33 (1993).

40. Bunin, G. Ecological communities with lotka-volterra dynamics. Physical Review E 95, 042414 (2017).

41. Fukami, T. Historical contingency in community assembly: integrating niches, species pools, and priority effects. Annual review of ecology, evolution, and systematics 46, 1–23 (2015).

42. Hu, J., Amor, D. R., Barbier, M., Bunin, G. – Gore, J. Emergent phases of ecological diversity and dynamics mapped in microcosms. Science 378, 85–89 (2022).

43. Sanchez, A. et al. Directed Evolution of Microbial Communities. Annual Review of Biophysics 50, 323–341 (2021). URL 10.1146/annurev-biophys-101220-072829.

44. Chang, C.-Y. et al. Engineering complex communities by directed evolution. Nature Ecology – Evolution 5, 1011–1023 (2021). URL http://www.nature.com/articles/s41559-021-01457-5.

45. Sanchez, A. et al. The community-function landscape of microbial consortia. Cell Systems 14, 122–134 (2023). URL https://www.sciencedirect.com/science/article/pii/S2405471222004999.

46. Weinreich, D. M., Lan, Y., Wylie, C. S. – Heckendorn, R. B. Should evolutionary geneticists worry about higher-order epistasis? Current Opinion in Genetics – Development 23, 700–707 (2013). URL https://www.sciencedirect.com/science/article/pii/S0959437X13001421.

47. Weinreich, D. M., Lan, Y., Jaffe, J. – Heckendorn, R. B. The Influence of Higher-Order Epistasis on Biological Fitness Landscape Topography. Journal of Statistical Physics 172, 208–225 (2018). URL http://link.springer.com/10.1007/s10955-018-1975-3.

48. Brookes, D. H., Aghazadeh, A. – Listgarten, J. On the sparsity of fitness functions and implications for learning. Proceedings of the National Academy of Sciences 119, e2109649118 (2022). URL https://pnas.org/doi/full/10.1073/pnas.2109649118.

49. Ballal, A. et al. Sparse Epistatic Patterns in the Evolution of Terpene Synthases. Molecular Biology and Evolution 37, 1907–1924 (2020). URL https://academic.oup.com/mbe/article/37/7/1907/5771371.

50. Sailer, Z. R. – Harms, M. J. Detecting High-Order Epistasis in Nonlinear Genotype-Phenotype Maps. Genetics 205, 1079–1088 (2017). URL https://academic.oup.com/genetics/article/205/3/1079/6066378.

51. Yang, G. et al. Higher-order epistasis shapes the fitness landscape of a xenobiotic-degrading enzyme. Nature Chemical Biology 15, 1120–1128 (2019). URL https://www.nature.com/articles/s41589-019-0386-3.

52. Poelwijk, F. J., Socolich, M. – Ranganathan, R. Learning the pattern of epistasis linking genotype and phenotype in a protein. Nature Communications 10, 4213 (2019). URL https://www.nature.com/articles/s41467-019-12130-8.

53. Hastie, T., Tibshirani, R. – Wainwright, M. Statistical Learning with Sparsity (Routledge, Boca Raton, 2015), 1st edition edn.

54. Candes, E. – Wakin, M. An Introduction To Compressive Sampling. IEEE Signal Processing Magazine 25, 21–30 (2008). URL http://ieeexplore. ieee.org/document/4472240/.

55. Chen, S. S., Donoho, D. L. – Saunders, M. A. suit. Atomic Decomposition by Basis Pur-SIAM Journal on Scientific Computing (2006). URL https://epubs.siam.org/doi/10.1137/S1064827596304010.

56. Poelwijk, F. J., Krishna, V. – Ranganathan, R. The Context-Dependence of Mutations: A Linkage of Formalisms. PLOS Computational Biology 12, e1004771 (2016). URL https://dx.plos.org/10.1371/journal.pcbi.1004771.

57. Yitbarek, S., Guittar, J., Knutie, S. A. – Ogbunugafor, C. B. Deconstructing taxa x taxa x environment interactions in the microbiota: A theoretical examination. bioRxiv 647156 (2021). URL https://www.biorxiv.org/content/10.1101/647156v2.

58. Doro, S. – Herman, M. A. On the Fourier transform of a quantitative trait: Implications for compressive sensing. Journal of Theoretical Biology 540, 110985 (2022). URL https://linkinghub.elsevier.com/retrieve/pii/S0022519321004057.

59. Marsland, R., Cui, W. – Mehta, P. A minimal model for microbial biodiversity can reproduce experimentally observed ecological patterns. Scientific Reports 10, 3308 (2020). URL https://www.nature.com/articles/s41598-020-60130-2.

60. Marsland, R., Cui, W., Goldford, J. – Mehta, P. The Community Simulator: A Python package for microbial ecology. PLOS ONE 15, e0230430 (2020). URL https://dx.plos.org/10.1371/journal.pone.0230430.

61. Nash, J. E. – Sutcliffe, J. V. River flow forecasting through conceptual models part I — A discussion of principles. of Hydrology 10, 282–290 (1970). https://www.sciencedirect.com/science/article/pii/0022169470902556.

62. Tibshirani, R. Regression Shrinkage and Selection Via the Lasso. Journal of the Royal Statistical Society: Series B (Methodological) 58, 267–288 (1996). URL https://onlinelibrary.wiley.com/doi/abs/10.1111/j.2517-6161.1996.tb02080.x.

63. Friedman, J., Higgins, L. M. – Gore, J. Community structure follows simple assembly rules in microbial microcosms. Nature Ecology – Evolution 1, 0109 (2017). URL http://www.nature.com/articles/s41559-017-0109.

64. Clark, R. L. et al. Design of synthetic human gut microbiome assembly and butyrate production. Nature Communications 12, 3254 (2021). URL http://www.nature.com/articles/s41467-021-22938-y.

65. Mehta, P. et al. A high-bias, low-variance introduction to Machine Learning for physicists. Physics Reports 810, 1–124 (2019). URL https://www.sciencedirect.com/science/article/pii/S0370157319300766.

66. Gould, A. L. et al. Data from: Microbiome interactions shape host fitness. Zenodo (2018). 10.5061/dryad.2sr6316.

67. Sanchez-Gorostiaga, A., Bajić, D., Osborne, M. L., Poyatos, J. F. – Sanchez, A. Data from: High-order interactions distort the functional landscape of microbial consortia. GitHub repository (2019). Https://github.com/djbajic/structure-function-bacilli.

68. Clark, R. L. et al. Data from: Design of synthetic human gut microbiome assembly and butyrate production. GitHub repository (2021). Https://github.com/RyanLincolnClark/DesignSyntheticGutMicrobiomeAssemblyFunction

69. Ye, J. C. Compressed sensing MRI: a review from signal processing perspective. BMC Biomedical Engineering 1, 8 (2019). URL 10.1186/s42490-019-0006-z.

70. Wohlberg, B. SPORCO: A Python package Journal URL for standard and convolutional sparse representations. Proceedings of the 16th Python in Science Conference 1–8 (2017). URL https://conference.scipy.org/proceedings/scipy2017/brendt_wohlberg.html. xConference Name: Proceedings of the 16th Python in Science Conference.

71. Boyd, S., Parikh, N. – Chu, E. Distributed Optimization and Statistical Learning Via the Alternating Direction Method of Multipliers (Now Publishers Inc, Hanover, MA, 2011).

72. Weinberger, E. D. Fourier and Taylor series on fitness landscapes. Biol. Cybern. 65, 321–330 (1991). URL https://doi.org/10.1007/BF00216965.

73. Pedregosa, F. et al. Scikit-learn: Machine Learning in Python. Journal of Machine Learning Research 12, 2825–2830 (2011). URL http://jmlr.org/ papers/v12/pedregosa11a.html.

74. Schmid, L., Gerharz, A., Groll, A. – Pauly, M. Machine Learning for Multi-Output Regression: When should a holistic multivariate approach be preferred over separate univariate ones? (2022). URL http://arxiv.org/abs/2201.05340. xArXiv:2201.05340 [cs, stat].

75. Friedman, J., Higgins, L. M. – Gore, J. Data and code from Friedman, Higgins, and Gore 2017 (Community structure follows simple assembly rules in microbial microcosms.” Nature ecology evolution 1.5 (2017): 0109) (2023). URL 10.5281/zenodo.8176044. 10.5281/zenodo.8176044.

76. Goldford, J. E. et al. Emergent simplicity in microbial community assembly. Science 361, 469–474 (2018). URL https://www.sciencemag.org/lookup/doi/10.1126/science.aat1168.

77. Gralka, M., Szabo, R., Stocker, R. – Cordero, O. X. Trophic Interactions and the Drivers of Microbial Community Assembly. Current Biology 30, R1176–R1188 (2020). URL https://linkinghub.elsevier.com/ retrieve/pii/S0960982220311611.

78. Estrela, S. et al. Functional attractors in microbial community assembly. Cell Systems S2405471221003793 (2021). URL https://linkinghub.elsevier.com/ retrieve/pii/S2405471221003793.

79. Enke, T. N. et al. Modular Assembly of Polysaccharide-Degrading Marine Microbial Communities. Current Biology 29, 1528–1535.e6 (2019). URL https://www.sciencedirect.com/science/article/pii/S0960982219303458.

80. Dal Bello, M., Lee, H., Goyal, A. – Gore, J. Resource–diversity relationships in bacterial communities reflect the network structure of microbial metabolism. Nature Ecology – Evolution 5, 1424–1434 (2021). URL https://www.nature.com/articles/s41559-021-01535-8.

81. O’Dwyer, J. P. Whence lotka-volterra? Theoretical Ecology 11, 441–452 (2018).

