## Supplementary Material for "Sparsity of higher-order landscape interactions enables learning and prediction for microbiomes"

### I. ECOLOGICAL LANDSCAPES AND THE WALSH-HADAMARD TRANSFORM

Given a pool of $S$ species, we consider all the $2^S$ ways of forming a seed community, given a fixed environment. Each combination can be denoted using a string of binary numbers, to indicate the initial presence/absence of species. The collection of these combinations or sub-communities is denoted as $\{\vec{\sigma}\}$. For $S = 3$ species in a pool, we have that, $\{\vec{\sigma}\} = \{000, 001, 010, 011, 100, 101, 110, 111\}$. There are $2^S = 2^3$ elements in this set, and each element can be denoted by the equivalent decimal representation of the string. For completeness, we include the ecologically-trivial case of all species being absent. For each species, we consider only a subset of $\{\vec{\sigma}\}$ since each species is initially present in only half of the sub-communities. We define this reduced space as $\{\vec{\sigma}_{(i)}\}$, for each species $i$. For species 1 in a pool with 3 species, this is $\{\vec{\sigma}_{(1)}\} = \{100, 101, 110, 111\}$

We define a mapping $\{\vec{\sigma}_{(i)}\} : \vec{a}_i$, where $a_i$ denotes the steady-state abundance vector of species indexed by $i$. We can define $S$ such maps, one for each species. These maps can be termed ecological landscapes [S1]:

$$f : \{0,1\}^{S-1} \to \mathbb{R} \tag{S1}$$

$\vec{\sigma}_{(i)}$ takes values in $\{0,1\}^{S-1}$ instead of $\{0,1\}^S$ since we assume the species $i$ is always present, at the start, in the relevant seed communities.

Consider a species in this pool, indexed by $i$, where we denote the abundance vector as $\vec{a}_i(\vec{\sigma})$, and coefficients of this vector, individually, as $N(\vec{\sigma})$. In the case of a 3-species pool, and for species 3, we have:

$$\begin{bmatrix} 001 \\ 011 \\ 101 \\ 111 \end{bmatrix} \mapsto \begin{bmatrix} N_{001} \\ N_{011} \\ N_{101} \\ N_{111} \end{bmatrix} \tag{S2}$$

We first consider a Walsh-Hadamard (WH) transform of this vector, without a diagonal matrix of weights pre-multiplying the transform. The matrix that implements this transform is defined recursively as:

$$H_{n+1} = \begin{bmatrix} H_n & H_n \\ H_n & -H_n \end{bmatrix} \text{with } H_0 = 1 \tag{S3}$$

These matrices are orthogonal s.t. $H_n H_n^T = 2^n \mathcal{I}_{2^n \times 2^n}$, symmetric s.t. $H_n = H_n^T$, and consist of $\pm 1$ as entries. Together, the symmetry and orthogonality imply that $H_n^{-1} = H_n/2^n$ [S2].

Multiplying the abundance vector with $H_n$ gives us the WH coefficients:

$$\vec{b}(\vec{\omega}) = H_n \vec{a}(\vec{\sigma})/2^n \tag{S4}$$

Here $n = S - 1$. The abundance vector can be computed, starting from the Walsh-Hadamard coefficients, using the relation: $\vec{a}(\vec{\sigma}) = H_n \vec{b}(\vec{\omega})$, where we used the fact that up to a factor $2^n$, the transform is equal to its inverse. To understand the coefficients $b$, which are a measure of landscape interactions, we look at the expansions of these coefficients in terms of abundance vector components, $N$. For clarity, we consider a case with $S = 4$, so that $n = S - 1 = 3$, and consider the abundance landscape of species 2. The Walsh-Hadamard coefficients are given by:

$$\begin{aligned}
b_0 &= b_{0\cdot 00} = (\textstyle\sum N_{[*\cdot **]})/8 \\
b_1 &= b_{0\cdot 01} = (\textstyle\sum N_{[*\cdot *0]} - N_{[*\cdot *1]})/8 \\
b_2 &= b_{0\cdot 10} = (\textstyle\sum N_{[*\cdot 0*]} - N_{[*\cdot 1*]})/8 \\
&\dots \\
b_6 &= b_{1\cdot 10} = (\textstyle\sum N_{[0\cdot 0*]} - N_{[0\cdot 1*]} - N_{[1\cdot 0*]} + N_{[1\cdot 1*]})/8 \\
b_7 &= b_{1\cdot 11} = (N_{0\cdot 00} - N_{0\cdot 01} - N_{0\cdot 10} + N_{1\cdot 01} - N_{1\cdot 00} + N_{1\cdot 01} + N_{1\cdot 10} - N_{1\cdot 11})/8
\end{aligned} \tag{S5}$$

where the $\cdot$ indicates that species 2 is always 1(present), and the symbol $*$ is a placeholder for 0 and 1, that is, we consider summing over both the values of 0 and 1 for these (background) species. The first coefficient $b_0$ then is simply the average abundance of species 2, considered over all the sub-communities. The coefficient $b_1 = b_{0101} = b_{0 \cdot 01}$ is a measure of the systematic effect of the presence and absence of species 4 on the abundance landscape of species 2, against the background of all other species (1 and 3) being present or absent. This is consistent with the symbol $*$ being in the places of species 1 and 3. $b_1$ is therefore, canonically, a pair-wise landscape interaction term, since it encodes the effect that species 4 has on the abundance of species 2. Similarly, $b_2$ is also a second-order term, which measures the effect of species 3 on the abundance landscape of species 2 (against the background of species 1 and 4 being present or absent). $b_6$ is a third-order term, which encodes the effect of species 1 and 3 on the abundance landscape of species 2. Finally, for $b_7$, there is no background to average over, and it is a measure of the systematic effect species 1,3, and 4 have on the abundance of species 2 in steady-state. In the main text, we use the transform $VH$, instead of simply $H$, with $V$ defined recursively as:

$$V_{n+1} = \begin{bmatrix} 0.5V_n & 0 \\ 0 & -V_n \end{bmatrix} \text{ with } V_0 = 1 \tag{S6}$$

$V$ is a diagonal weighting matrix that takes into account the order of interactions, to account for averaging over different numbers of terms as a function of the order of interactions [S3–S5]. In our case, inclusion of $V$ is an ecologically motivated choice that down-weighs higher-order landscape interactions. This is based on the assumption that while higher-order landscape interactions are allowed, lower order interactions are relatively more likely, and the weighting encoded in $V$ captures this expectation. In terms of unweighted Walsh-Hadamard coefficients, we can define the new coefficients, $\beta$, as

$$\beta_j = b_j \cdot (-1)^{q_j} \cdot 2^{q_j} \tag{S7}$$

where $q_j$ is the order of the coefficient, which is simply the sum of the number of 1s in the index of the $b$ coefficient (excluding the 1 corresponding to the focal species). The second-order weighted-WH coefficient, $\beta^{(2)}$, is given by:

$$\beta^{(2)} = \beta_{0 \cdot 01} = -\frac{1}{4}\left[\left(N_{0100} - N_{0101}\right) + \left(N_{0110} - N_{0111}\right) + \left(N_{1100} - N_{1101}\right) + \left(N_{1110} - N_{1111}\right)\right]$$

And the fourth-order weighted-Walsh-Hadamard coefficient is given by:

$$\beta^{(4)} = \beta_{1 \cdot 11} = -\left[N_{0100} - N_{0101} - N_{0110} + N_{0111} - N_{1100} + N_{1101} + N_{1110} - N_{1111}\right]$$

#### A. Explanatory power of the Walsh-Hadamard coefficients

We suppose that we have access to the entire combinatorial data-set for a microbiome, with all the $\vec{a}$ and $\vec{b}$ components known. Both these vectors contain the same information, although we may observe only the abundances. However, these vectors are large, with $\vec{a}$ consisting of potentially $2^{S-1}$ non-zero coefficients. Even a modest increment in $S$ means sampling a lot of experimental data, which become impossible as $S$ increases. We instead explore the distribution of information in the Walsh-Hadamard coefficients, $\vec{b}$.

To this effect, we find the top $k$ coefficients of $\vec{b}$ which are able to explain most of the variance in $\vec{a}$. We construct the vector $\vec{b}^{(k)}$, with $k$ coefficients chosen from $\vec{b}$ and remaining $2^n - k$ coefficients set to 0. Here we set $S - 1 = n$. For a given $k$, the vector $\vec{b}^{(k)}$ with the highest explanatory power will minimize the distance: $\|\vec{a} - \vec{a}^{(\text{approx.})}\|^2$, where $\vec{a}$ is the original abundance vector which is also equal to $H_n\vec{b}(\omega)$, and $\vec{a}^{(\text{approx.})} = H_n\vec{b}^{(k)}(\omega)$. Because of the orthogonality and symmetry of the unweighted-WH transform, the minimization problem is equivalent in both the representations, i.e :

$$\|\vec{a} - \vec{a}^{(\text{approx.})}\|^2 = 2^n\|\vec{b} - \vec{b}^{(k)}\|^2 \tag{S8}$$

For a given $k$, the vector $\vec{b}^{(k)}$ which is a solution to Equation S8 is also called the best sub-setting approximation [S6]. In this case, there exists a simple recipe to find the $k$ most significant coefficients—we need only to sort the $b$ coefficients by absolute values and keep the top $k$ coefficients. This implies we can easily calculate the variance explained by $k$ coefficients as $100 \times \text{PS}(\vec{a}, \vec{a}^{(\text{approx.})})$ where

$$\text{PS} = \frac{1}{2 - R^2(\vec{a}, \vec{a}^{(\text{approx.})})} \tag{S9}$$

$R^2$ is the coefficient of determination. However, in the main text, we use the transform $VH$, instead of simply $H$, with $V$ defined recursively as:

$$V_{n+1} = \begin{bmatrix} 0.5V_n & 0 \\ 0 & -V_n \end{bmatrix} \text{with } V_0 = 1 \tag{S10}$$

In terms of un-weighted Walsh-Hadamard coefficients, we can define the new coefficients as

$$\beta_j = b_j \cdot (-1)^{q_j} \cdot 2^{q_j} \tag{S11}$$

where $q_j$ is the order of the coefficient, which is simply the sum of the number of 1s in the index of the $b$ coefficient (excluding 1 corresponding to the focal species). For the case of 4 species considered above, this means $\beta_0 = b_0$, $\beta_1 = (-1)^1 \cdot 2^1 \cdot b_1$, and $\beta_7 = (-1)^3 \cdot 2^3 b_7$. It remains easy to find the best sub-setting approximation in this case: squared error in the abundance space is now equivalent to weighted squares in the $\beta$ coefficients:

$$\|\vec{a} - \vec{a}^{(approx.)}\|^2 = \sum_j u_j^2 (\beta_j - \beta_j^{best})^2 \tag{S12}$$

with $u_j = 2^{\frac{n}{2} - q_j}$. This means that to find the top $k$ coefficients we need to sort the absolute values of the $\beta$ coefficients, while taking into account the order of the coefficient, as prescribed by $u_j$. However, since $\beta_j = b_j \cdot (-1)^{q_j} \cdot 2^{q_j}$, sorting $u_j \beta_j$ by absolute value is the same as sorting $b_j$ by absolute values, and we get exactly the same explained variance for the top $k$ coefficients in both the bases.

#### B. Explained variance for Walsh-Hadamard and weighted Walsh-Hadamard coefficients

For the in silico 16-species datasets, we consider the variance explained by different terms in the Walsh-Hadamard and the weighted Walsh-Hadamard bases. By considering the sub-setting approximations as described in the section above, we find, as expected, that the variance explained by the $b$ and the $\beta$ coefficients is the same when we consider the same number of top $k$ coefficients. We demonstrate this equivalence, for two different species, chosen from the 16-species in silico community, in Fig. S1 panels A and B, respectively.

We refer the reader to [S2–S6] for an excellent treatment of Walsh-Hadamard matrices in the context of evolutionary biology and statistical genetics.

### II. SPARSITY OF HIGHER-ORDER LANDSCAPE INTERACTIONS IN AN EXPERIMENTAL DATA-SET

Gould *et al.* [S7, S8] sampled a combinatorially-complete landscape of 5-species. This data-set proves to an excellent test-bed for exploring the concept of sparsity that we introduce and study in this work. It also brings to the fore the notion of sparsity of higher-order landscape interactions. Figure S2 shows that all species in this particular system survive at steady-state, given that they were present at the start in the seed community: all steady-states in this data-set are maximally diverse, with the same diversity as the seed community. In Figure S2, each sub-community is labeled and identified by the unique combination of whether each species is initially present or absent. In this representation, there is no evident sparsity or other simplifying feature; sampling a species' abundance in a few sub-communities does not lead to any obvious way to predict its abundance in other sub-communities.

On the other hand, we can calculate the weighted Walsh-Hadamard (WH) coefficients corresponding to the relative abundance vector of one species, A. *orientalis*, and we summarize the results in Figure S3A. We show the size of these coefficients in Figure S3A, as a function of their order. i.e. order 2 indicates pairwise dependencies between A. *orientalis* and one other species, while higher orders indicate the dependence of A. *orientalis*' abundance on combinations of two or more other species. We see that there are only a handful of large coefficients (**indicating sparsity**), and also a bias towards lower order interactions (**indicating the sparsity of higher-order interactions**). Figure S3B shows that most of the variance in A. *orientalis* abundances can be explained by the first three weighted-WH coefficients, all of which are low-order. In summary, this system with maximally-diverse steady-states has a sparse representation with few higher-order interactions, and using only the largest coefficients in the Walsh-Hadamard transform works well in terms of reconstructing community composition.

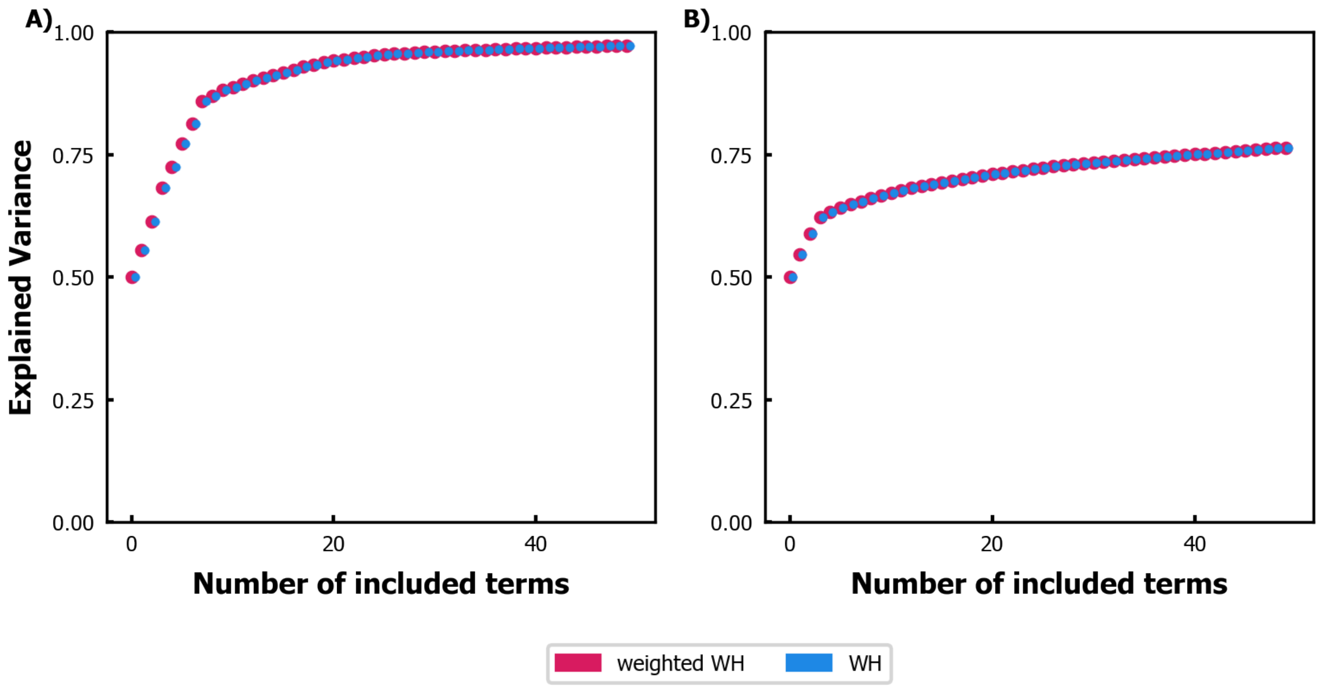


FIG. S1: **Variance explained by top $k$ coefficients using the weighted and unweighted Walsh-Hadamard transform.** The variance explained by top 50 coefficients is the same in both weighted Walsh-Hadamard transform (shown in red) and the unweighted Walsh-Hadamard transform (points shown in blue), when considering two randomly selected species from the in silico community. Panel **A)** shows the comparison for one species, while panel **B)** shows the comparison for a second species. Note that the computation of explained variance in these cases required access to the entire abundance datasets.

### III. THE MICROBIAL CONSUMER RESOURCE MODEL

To generate a combinatorially-complete data-set of species abundances at steady-state, we considered an in silico community of 16 species in well-mixed, chemostat conditions. The dynamics of this community is described by the microbial consumer resource model (MiCRM) [S9–S11].The MiCRM considers both the species and resource dynamics explicitly. For $S$ species and $M$ resources, the matrix, $C_{i\alpha}^{S\times M}$, consists of elements, $c_{i\alpha}$, where each element is the uptake rate for resource $\alpha$ by species $i$. Interaction between the species is mediated by resource consumption which gives rise to competitive interactions, but also incorporates cross-feeding terms, by allowing for species to leak a fraction, $l$, of the resources it consumes in the form of metabolic byproducts. The composition of these byproducts is specified by the metabolic leakage matrix $\mathcal{D}$, with matrix element $\mathcal{D}_{\alpha\beta}$ specifying the amount of resource $\beta$ leaked when the species consumes resource $\alpha$. Each row of the leakage matrix sums to one. The MiCRM is defined by the following equations, where $n_i$ are the species abundances and $Y_\alpha$ are the resource concentrations:

$$\begin{aligned}\frac{dn_i}{dt} &= g_i n_i \left[(1-l)\sum_{\alpha=1}^{M} w_\alpha \sigma\left(c_{i\alpha}Y_\alpha\right) - m_i\right], \\ \frac{dY_\alpha}{dt} &= \tau^{-1}\left(Y_\alpha^0 - Y_\alpha\right) - \sum_{i=1}^{S} n_i \sigma\left(c_{i\alpha}Y_\alpha\right) + l\sum_{i=1}^{S}\sum_{\beta=1}^{M} n_i \mathcal{D}_{\alpha\beta}\frac{w_\beta}{w_\alpha}\sigma\left(c_{i\beta}Y_\beta\right).\end{aligned} \tag{S13}$$

We chose the resource uptake function to have a linear form: $\sigma(x) = x$. We supplied one resource externally, at a supply rate of $Y_{\alpha=5}^0 = 100$. Since cross-feeding is allowed, there are in total 20 metabolites to go around in this community. The leakage fraction, that allows for this cross-feeding, was set to $l = 0.6$ for all species. The leaked fraction of the consumed resource can take up many forms, and this is prescribed by the matrix $\mathcal{D}$. We sampled each column of this matrix from a Dirichlet distribution with sparsity parameter 0.3. The growth rate, $g_i$, and the maintenance rate, or the minimal energy uptake for maintenance of species $i$, $m_i$, were both set to 1. To reflect chemostat conditions, timescale of resource dilution, $\tau$ was also set to 1, as were the resource qualities $w$. We sampled

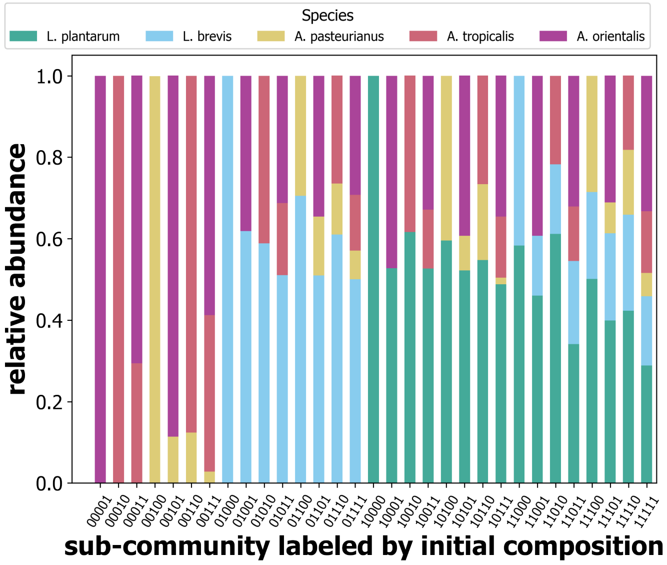


FIG. S2: Relative abundances of species in different assemblages in the fly gut data-set [S7, S8]. Each sub-community is labeled and identified on the x-axis by the unique combination of whether each species is initially present or absent. In this data-set, all steady-states are maximally diverse, with the same diversity as the seed community, and no obvious sparsity.

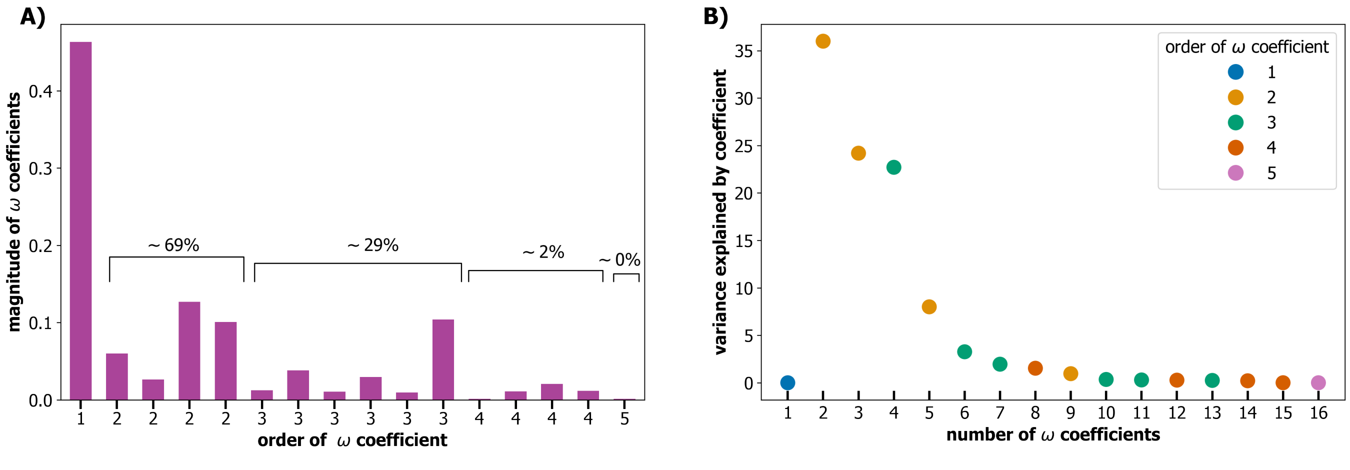


FIG. S3: For the abundance landscape of species *A. orientalis*, the Walsh-Hadamard representation is a sparse one, with most of the variance explained by lower-order coefficients. Around 70% of the variance in the abundance can be explained by the first four coefficients, all of which are low-order.

the uptake rates $c_{i\alpha}$ from a binary-gamma distribution using a prescription detailed in Ref. [S12]. We chose the mean, $\mu_c = \langle C_i \rangle = 15.0$, and $\sigma_c^2 = \text{var}(C_i) = 2.3$.

### IV. PREDICTION ABILITY IS RELATED TO THE VARIANCE EXPLAINED BY LOWER-ORDER LANDSCAPE INTERACTIONS

For the simulated data-set, where we have access to all the data, we consider the distribution of variance explained by different $\beta$ terms, across orders of landscape interactions. We define a measure, SC:

$$SC = \sum_i f_i((S-1)/2 - i) \tag{S14}$$

where $f_i$ is the average variance explained by all the $\beta$ terms at order $i$, and $S$ is the number of species in the pool. This measure penalizes species whose abundance landscape is not dominated by lower-order terms by allowing negative terms in the sum in Equation S14, and reducing the subsequent $SC$ value. We expect species that are dominated by lower-order landscape interactions to have higher $SC$ values, and that their abundance landscape will be easier to learn from limited data by compressive sensing algorithms. To probe this correlation, we calculate the Spearman's rank correlation coefficient, $\rho$, between the theoretically calculated $SC$ value and the prediction score on out-of-sample points from compressive sensing when only 1% of the data is available for training. The association is plotted in Fig. S4A, with Spearman's rank coefficient of 0.97, which is in line with our expectation of prevalent lower-order landscape interactions for easier-to-predict species.

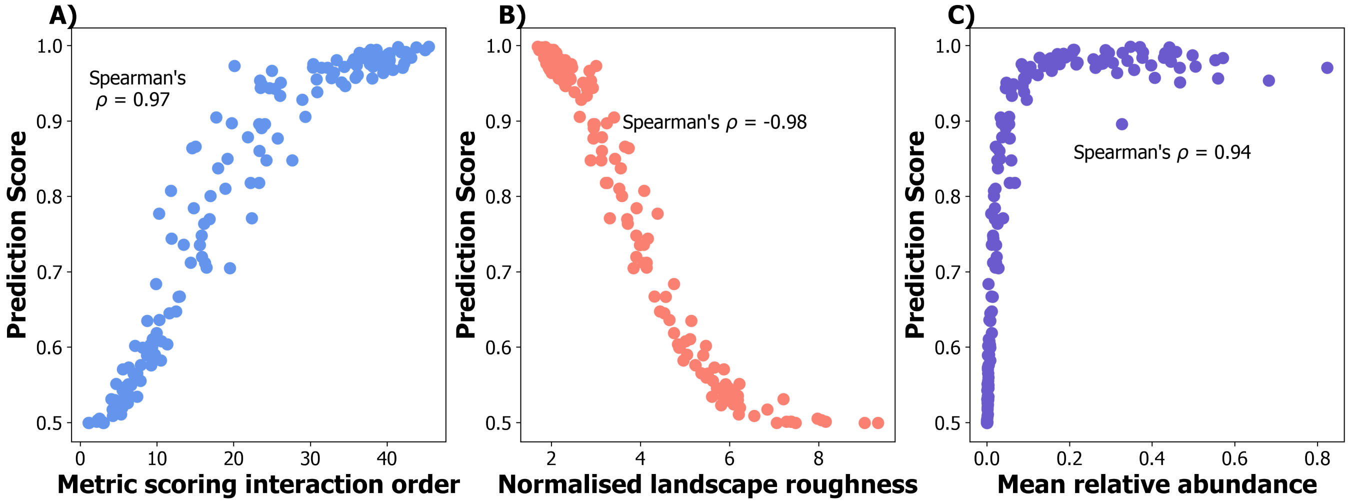


FIG. S4: **Sparsity of higher-order landscape interactions and higher relative mean abundance are associated with higher prediction scores. A)** For the simulated data-set with 16 species, and across 10 simulation replicates, we computed a value which measures the importance of lower-order interactions in terms of fraction of explained variance using equation S14. The large value of the Spearman's rank correlation coefficient demonstrates that species with significant lower-order interactions are predicted well from limited data using compressive sensing algorithms. **B)** Species which have a smoother abundance landscape are also predicted better. Smoothness of the landscape is defined in terms of lower relative roughness, as computed using equation S17. **C)** Species with higher mean relative abundance are associated with better prediction scores. Prediction score was calculated using Equation 2 of the main text on out-of-sample predicted abundances. Prediction scores (at optimal regularization parameters) are reported for simulated data-sets with 1% training data, and averaged across 5 runs.

It is also possible to define a measure of how rugged the abundance landscape of a species is. For this, as in Ref. [S2], we may define a trait roughness, $\mathcal{R}(\vec{a}_i)$, which is a sum of the "influence" of all other species on the abundance of species $i$. In terms of the unweighted Walsh-Hadamard transformed coefficients, $b$, this is simply:

$$\mathcal{R}(\vec{a}_i) = \sum_{k=0}^{2^{S-1}} (b_i)_k^2 \tag{S15}$$

In terms of the weighted Walsh-Hadamard transformed coefficients, $\beta$, the roughness measure can be defined as :

$$\mathcal{R}(\vec{a}_i) = \sum_{k=0}^{2^{S-1}} h_k(q)(\beta_i)_k^2 \tag{S16}$$

where $h_k(q) = \frac{1}{2^{2(q+1)-S}}$, $q$ is the sum of the number of 1s in the index of the $\beta$ coefficient, and therefore $h_k(q)$ is an order-dependent term. For every species, we may define a relative roughness value, $\imath$, using the variance in the abundance of a species, $\sigma^2(\vec{a}_i)$, :

$$\imath(\vec{a}_i) = \mathcal{R}(\vec{a}_i)/\sigma^2(\vec{a}_i). \tag{S17}$$

For the simulated data-set, we computed the relative roughness for all species in all the 10 simulation pools, and looked at the correlation between prediction scores on out-of-sample data (for 1% training data) and the relative roughness. This strong correlation is plotted in Fig. S4B, where the large negative Spearman's rank correlation coefficients indicates that species with higher relative roughness are not predicted well using limited data. Furthermore, the mean relative abundance of a species, considered over all the sub-communities, is also associated with higher prediction scores, as shown in S4C.

### V. COMPRESSIVE SENSING WITH UNWEIGHTED WALSH-HADAMARD TRANSFORM

In the main section, we used a weighted Walsh-Hadamard matrix, $\Omega = VH$, where we implement an ecologically motivated choice that down-weights higher-order landscape interactions and biases inference towards lower-order interaction terms. This weighing is encoded in the diagonal matrix $V$, which consists of landscape interaction-dependent factors. In Figs. S5 and S6, we show that using compressive sensing with our transform of choice does better on both the simulated and real data-sets than an unweighted Walsh-Hadamard transform.

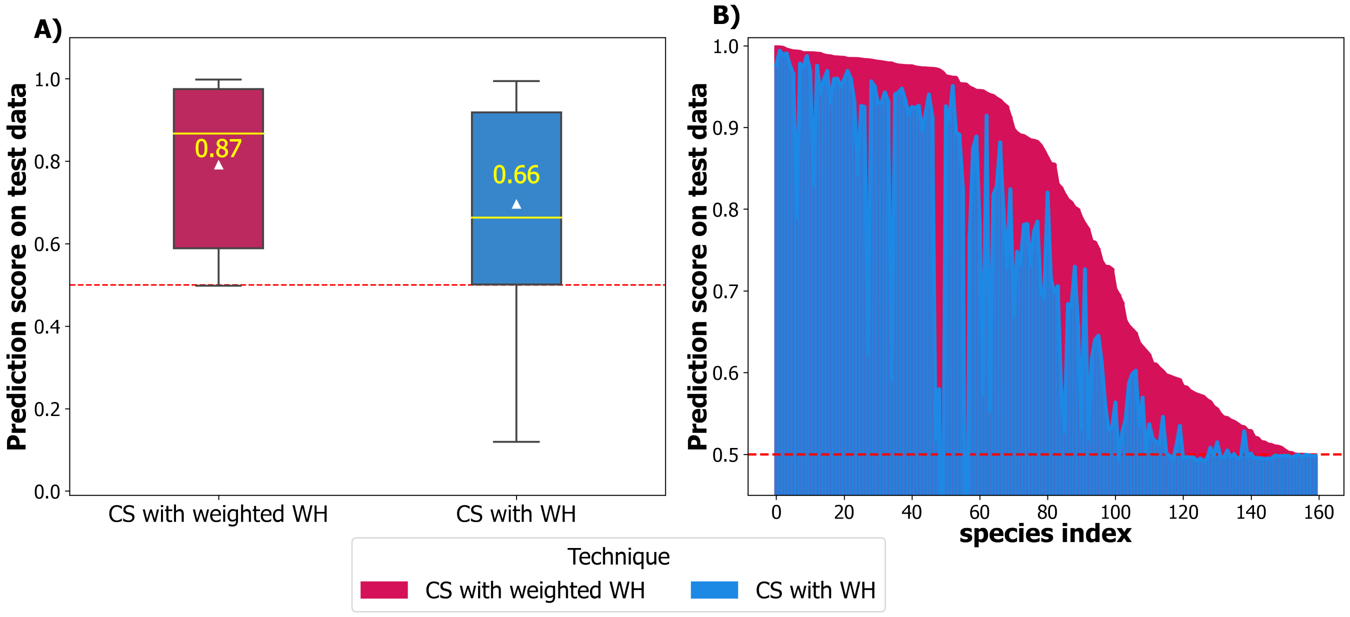


FIG. S5: **Compressive sensing with a weighted WH transform predicts better than an unweighted transform on the simulated data-set. A)** For the in silico community with 16 species, and across 10 pools of simulations, we found that compressive sensing with a weighted Walsh-Hadamard transform, where the weights are chosen so as to down-weight higher-order landscape interaction terms, does significantly better than an unweighted Walsh-Hadamard transform. Box-plots show that median values of out-of-sample prediction score for compressive sensing with weighted and unweighted WH transforms are 0.87 and 0.66 respectively. Permutation test p-value of 0.016 for difference in the means indicates that this difference in performance is significant. **B)** A species-by-species matched comparison for each of the 160 species (16 species in 10 simulation pools) shows that CS with weighted WH transform leads to better prediction scores.

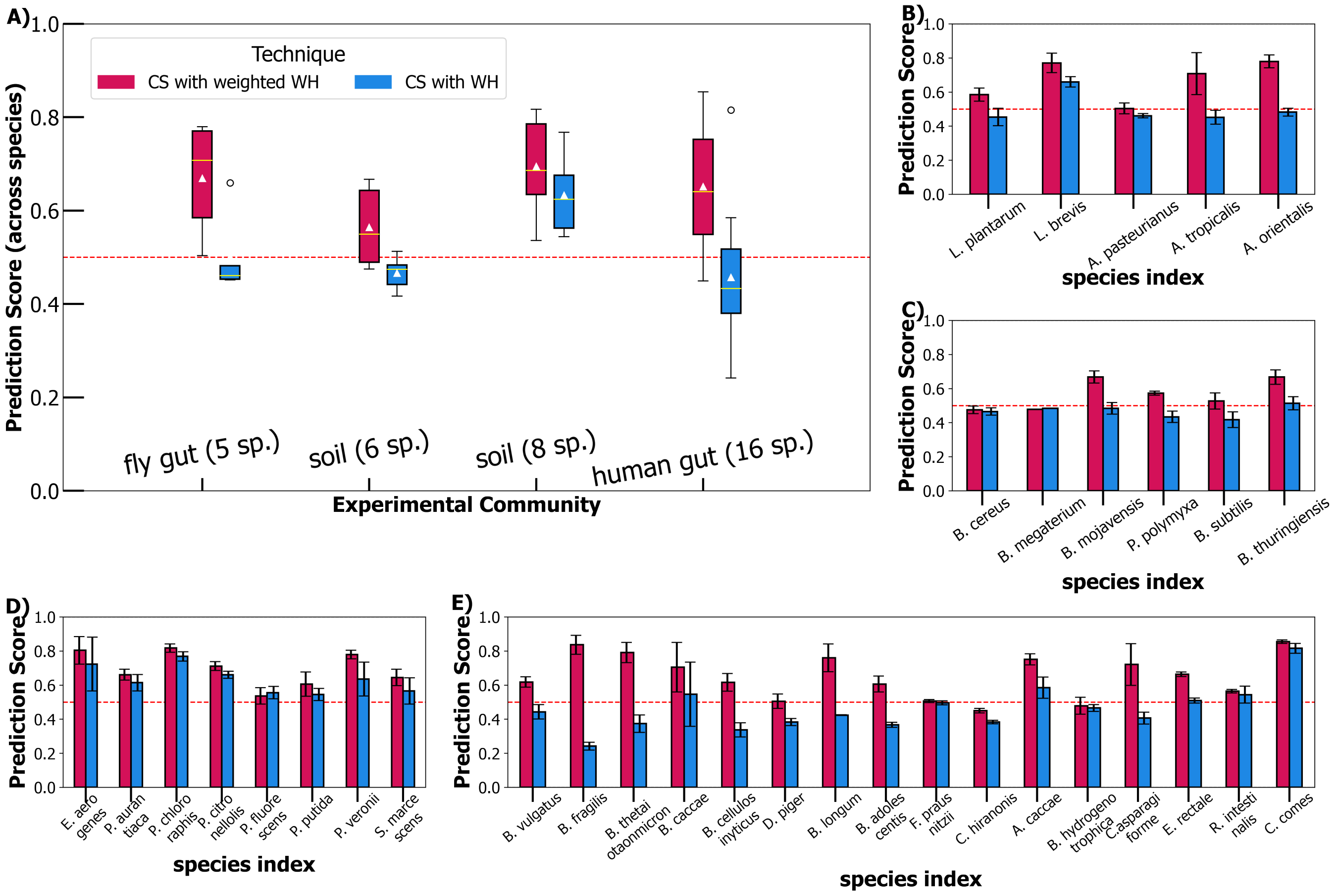


FIG. S6: **Compressive sensing with a weighted WH transform predicts better than an unweighted transform on experimental data. A)** For the 4 experimental data-sets considered in this paper, we found that compressive sensing with a weighted Walsh-Hadamard transform, where the weights are chosen so as to down-weight higher-order interaction terms, does significantly better than a plain, unweighted Walsh-Hadamard transform. Box plots show that median values of out-of-sample prediction score for compressive sensing with weighted transform are consistently higher, indicating better performance at a community level. **B)** For the 5-species data-set in Ref. [S7, S8], we found that a weighted WH transform is a better choice for predicting abundances using compressive sensing. A permutation test p-value indicates that the difference in means is significant with p-value .03. For datasets in Refs. [S13–S17], we demonstrate that using the weighted transform is a better choice as shown in a matched comparison at a species level in panels **C)**, **D)**, and **E)**. Permutation test p-values are .031, .008, and $1.52 \times 10^{-5}$ respectively.

We found, by analyzing the contribution of the $\beta$ coefficients at different orders, that lower-order coefficients contribute disproportionately to the explained variance. This is the case when considering the explained variance vs. order of significant terms in the unweighted Walsh-Hadamard transform as well, as shown in Fig. S1, when considering all the data for the in silico communities. This happens to be the case in a real community as well, which we describe above, in Section 2. We can incorporate this finding when trying to infer abundances from limited data. When inferring the $\vec{\beta}$ or the $\vec{b}$ vectors, which are expected to be sparse, with the sparsity coming from relative rarity of significant higher-order terms, we can condition the sparse recovery algorithm to heavily penalize the $b$ coefficients that correspond to higher-order landscape interactions. This can be implemented in the LASSO/BPDN (LASSO : Least Absolute Shrinkage and Selection Operator, BPDN : Basis Pursuit de-Noising) algorithms of compressive sensing by a weighted penalty, $\vec{\lambda}$, instead of an indentical parameter, $\lambda$, for each $b_j$. In the case of microbial data-sets, this means $\lambda_j$ corresponding to lower-order $b_j$ must be smaller when compared to $\lambda_j$ for higher-order $b_j$. This can be accommodated in the LASSO/BPDN algorithm:

$$\underset{b}{\operatorname{argmin}}\Big(\frac{1}{2}||Db - a||_2^2 + \lambda||b||_1\Big) \tag{S18}$$

Here, $D$ is the partial inverse (unweighted) Walsh-Hadamard matrix with rows sampled according to available observed abundances. When incorporating different $\lambda$s for different coefficients, we may write this as:

$$\underset{b}{\text{argmin}}\Big(\frac{1}{2}||Db - a||_2^2 + ||\Lambda b||_1\Big) \quad \text{(S19)}$$

Here $\Lambda$ is a diagonal matrix with entries $\lambda_j$. Implementing this order-weighted penalty matrix can be done in the Python package [S18] we use. Another way of implementing an order-weighted regularization is to make changes to the $D$ matrix itself. In this case, the optimization problem reduces to :

$$\underset{b}{\text{argmin}}\Big(\frac{1}{2}||D\Lambda^{-1}\beta - a||_2^2 + ||\beta||_1\Big) \quad \text{(S20)}$$

Here $\beta = \Lambda b$, which is proportional to the transformation we have in Equation S11. Thus, compressive sensing with the weighted transform, $D\Lambda^{-1}$, with $\Lambda = V$, is equivalent to heavily penalizing higher-order coefficients in the original LASSO/BPDN program. In Fig. S7, we include comparisons of prediction scores for out-of-sample abundances using

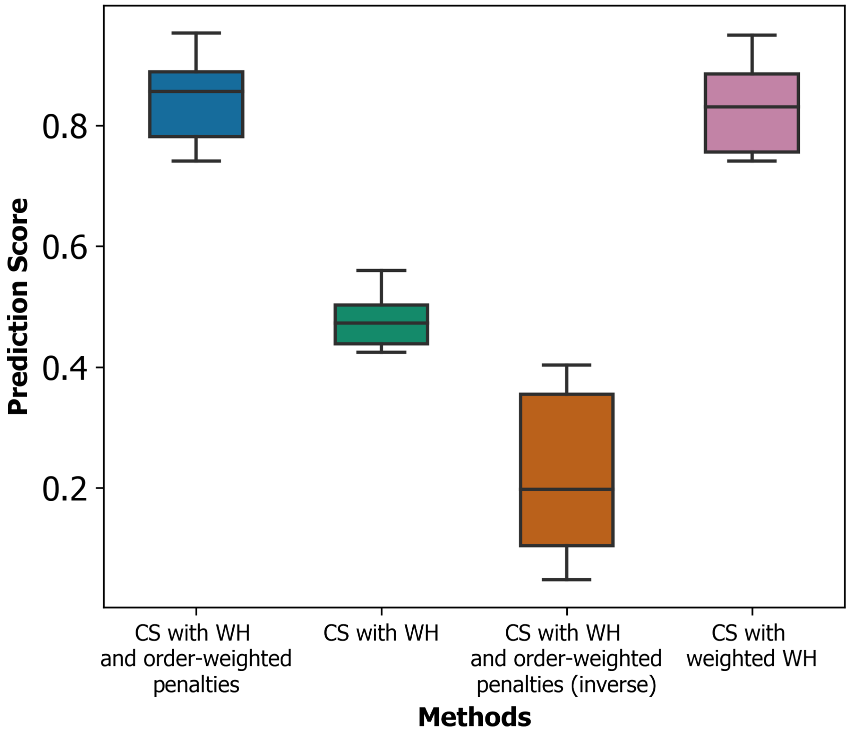


FIG. S7: **Compressive sensing with a weighted WH transform is equivalent to compressive sensing with an unweighted transform with order-weighted regularization parameters.** The algorithm we used to implement compressive sensing is called BPDN/LASSO [S19, S20], and is a convex optimization problem as given in equation S18. To recover sparse coefficients $b$, we need to choose a design matrix, $D$, and regularization terms $\lambda$, that penalize the $l_1$ norm of the sparse vector. We demonstrate that the weighted transform used throughout the main text and whose performance is shown here in the box-plot labelled "CS with weighted WH" is comparable to "CS with WH and order-weighted penalties". This means that changing the design matrix from $H^{-1}$ to $(VH)^{-1}$ is the same as imposing order-weighted penalties, where the ordering is proportional to the diagonal elements of $V$. This translates to heavily penalizing higher-order terms in the sparse vector. Both of these choices in LASSO/BPDN perform better than LASSO/BPDN with unweighted WH transform and unweighted penalties, as shown in the box-plot labelled "CS with WH". It is also possible to then choose the penalty/regularization terms, $\lambda_j$, to be systematically different from what we expect, which is to heavily penalize lower-order coefficients. This leads to an expected degradation in prediction score, as seen in "CS with WH and order-weighted penalties (inverse)" which specifically does worse than "CS with WH". For this figure, we chose a simulated 10-species in silico community, drawn from the 16-species in silico community used elsewhere. Predictions are reported on out-of-sample abundances with 5% data available for training.

4 different methods: we implement equation S19, with order-weighted penalties such that higher-order terms are penalized more, with the weighing given by the diagonal elements of $V$. The matrix $D$, the design matrix, is a partial

inverse unweighted Walsh-Hadamard matrix. The performance of compressive sensing, using this method, is shown in left-most box-plot (blue color) labelled "CS with WH and order-weighted penalties". Note that the performance for this case is comparable to the performance for the method: "CS with weighted WH", where we use equation S20. This second case (results shown in pink box-plot) is the algorithm we use in the main text, with a transform $VH$. Both of these implementations do better than LASSO/BPDN as implemented using equation S18 where the penalties are agnostic to the coefficients and the transform is an unweighted Walsh-Hadamard one. This is seen in Fig. S7, in the (green-colored) box-plot labelled "CS with WH". Along these lines, there exists a way to worsen the performance of LASSO/BPDN by choosing order-weighted penalties, where the weights are inversely chosen, such that lower-order terms are penalized more. This degradation in performance is expected in the microbial data-sets, and is demonstrated in the box-plot labelled: "CS with WH and order-weighted penalties (inverse)".

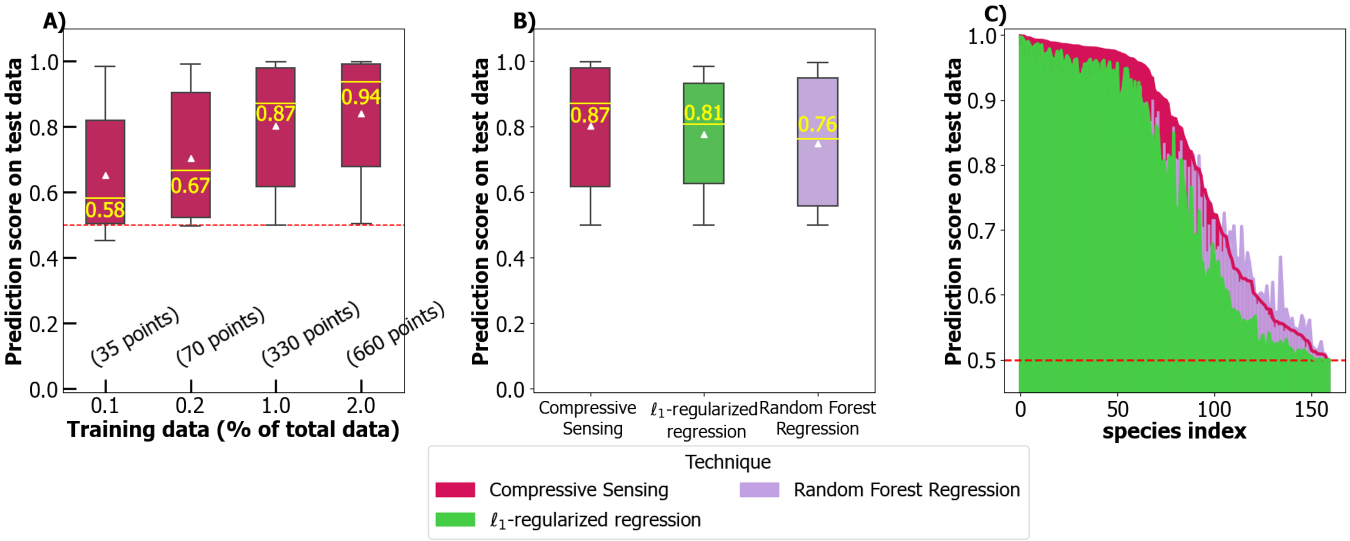


FIG. S8: **Compressive sensing with a weighted Walsh-Hadamard transform does well on simulated datasets when the data consists of absolute abundances.** Compared to $l_1$-regularized regressions and random forest regression, predictions of compressive sensing on simulated data-set continue to be more accurate, as shown in panel **B)**, where the median values of prediction scores are higher. A matched comparison at the species-level in **C)** demonstrates the accuracy of this method.

### VI. PREDICTIONS ON ABSOLUTE ABUNDANCES

In the main text, in Figures 3 and 4, we work with relative abundances. Here, we also compare the performance of compressive sensing on absolute abundances. All the experimental datasets had a measure of absolute abundances, with Gould et al. and Sanchez-Gorostiaga et al. reporting colony-forming units (CFUs) [S7, S13], and Friedman et al. and Clark et al. reporting $OD_{600}$ measurements [S15, S16]. In the simulated data-set, we generate cell count steady-state abundances. The results for compressive sensing performance are shown in Figs S8 and S9.

### VII. SELECTION OF THE 16-SPECIES POOL FROM THE 25-SPECIES COMMUNITY STUDIED BY CLARK ET. AL [S16]

As outlined in the Methods section, for this data-set, we considered a sub-set of 16 species, with the other 9 always being absent. Since there are many possible sub-sets of 16 species in the background of some consistently-absent 9 species, we selected a 16-species pool such that the number of experimental data-points available to us was maximized. The distribution of available data-points with different number of species consistently absent is shown in Figure S10.

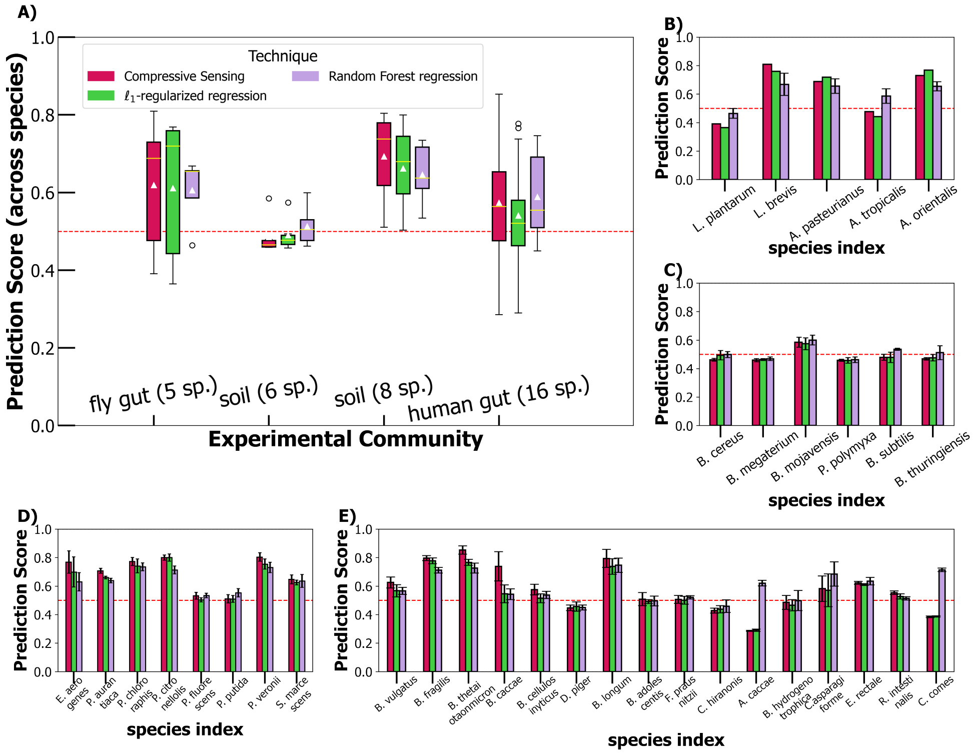


FIG. S9: **Compressive sensing with a weighted Walsh-Hadamard transform does well on experimental data when the data consists of absolute abundances.** Predictions of compressive sensing continue to be accurate on the two largest real microbiomes, as can be seen from the median values (horizontal yellow bars) of the box plots in panel **A)**.

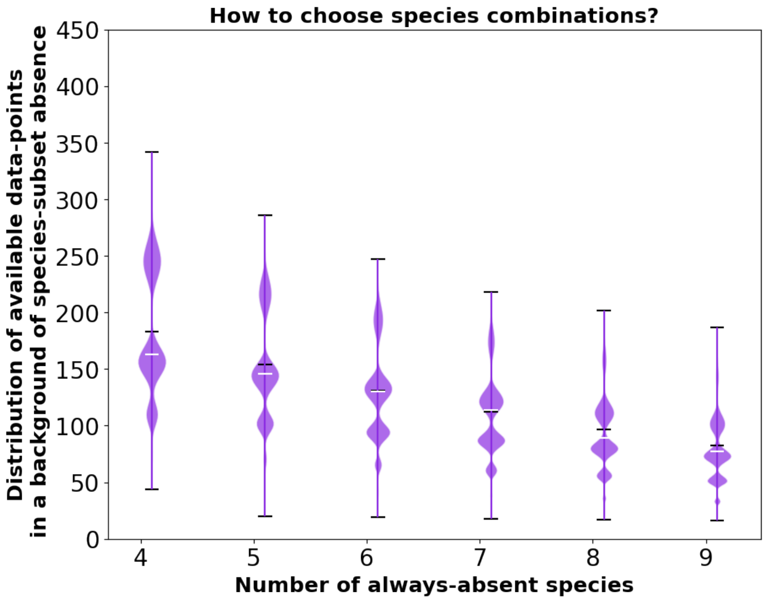


FIG. S10: For the data-set in Ref. [S16] with 25 species, we chose a sub-set of 16 species in the background of 9 always-absent species. This sub-set selection was made after noting the distribution of available experimental combinations. Although the most number of data-points ($\sim$ 350) are available for a set of 21 species with 4 always-absent species, computational complexity of the compressive sensing pipeline led us to choose a 16-species subset, with this particular sub-set giving us 187 experimental combinations, from a theoretical possible $2^{16}-1$ combinations.

**Bibliography**

---

[S1] A. B. George and K. S. Korolev, PLOS Computational Biology **19**, e1010570 (2023), ISSN 1553-7358, URL https://journals.plos.org/ploscompbiol/article?id=10.1371/journal.pcbi.1010570.
[S2] S. Doro and M. A. Herman, Journal of Theoretical Biology **540**, 110985 (2022), ISSN 00225193, URL https://linkinghub.elsevier.com/retrieve/pii/S0022519321004057.
[S3] D. M. Weinreich, Y. Lan, C. S. Wylie, and R. B. Heckendorn, Current Opinion in Genetics & Development **23**, 700 (2013), ISSN 0959-437X, URL https://www.sciencedirect.com/science/article/pii/S0959437X13001421.
[S4] F. J. Poelwijk, V. Krishna, and R. Ranganathan, PLOS Computational Biology **12**, e1004771 (2016), ISSN 1553-7358, URL https://dx.plos.org/10.1371/journal.pcbi.1004771.
[S5] F. J. Poelwijk, M. Socolich, and R. Ranganathan, Nature Communications **10**, 4213 (2019), ISSN 2041-1723, URL https://www.nature.com/articles/s41467-019-12130-8.
[S6] D. M. Weinreich, Y. Lan, J. Jaffe, and R. B. Heckendorn, Journal of Statistical Physics **172**, 208 (2018), ISSN 0022-4715, 1572-9613, URL http://link.springer.com/10.1007/s10955-018-1975-3.
[S7] A. L. Gould, V. Zhang, L. Lamberti, E. W. Jones, B. Obadia, N. Korasidis, A. Gavryushkin, J. M. Carlson, N. Beerenwinkel, and W. B. Ludington, Proceedings of the National Academy of Sciences **115**, E11951 (2018), ISSN 0027-8424, 1091-6490, URL http://www.pnas.org/lookup/doi/10.1073/pnas.1809349115.
[S8] A. L. Gould, L. Zhang, Vivian andLamberti, E. W. Jones, B. Obadia, N. Korasidis, A. Gavryushkin, J. M. Carlson, N. Beerenwinkel, and W. B. Ludington, Zenodo (2018), https://doi.org/10.5061/dryad.2sr6316.
[S9] R. Marsland, W. Cui, and P. Mehta, Scientific Reports **10**, 3308 (2020), ISSN 2045-2322, URL https://www.nature.com/articles/s41598-020-60130-2.
[S10] R. MacArthur, Theoretical Population Biology **1**, 1 (1970), ISSN 00405809, URL https://linkinghub.elsevier.com/retrieve/pii/0040580970900390.
[S11] R. Marsland, W. Cui, J. Goldford, and P. Mehta, PLOS ONE **15**, e0230430 (2020), ISSN 1932-6203, URL https://dx.plos.org/10.1371/journal.pone.0230430.
[S12] C.-Y. Chang, J. C. C. Vila, M. Bender, R. Li, M. C. Mankowski, M. Bassette, J. Borden, S. Golfier, P. G. L. Sanchez, R. Waymack, et al., Nature Ecology & Evolution **5**, 1011 (2021), ISSN 2397-334X, URL http://www.nature.com/articles/s41559-021-01457-5.
[S13] A. Sanchez-Gorostiaga, D. Bajić, M. L. Osborne, J. F. Poyatos, and A. Sanchez, PLOS Biology **17**, e3000550 (2019), ISSN 1545-7885, URL https://dx.plos.org/10.1371/journal.pbio.3000550.
[S14] A. Sanchez-Gorostiaga, D. Bajić, M. L. Osborne, J. F. Poyatos, and A. Sanchez, GitHub repository (2019), https://github.com/djbajic/structure-function-bacilli.
[S15] J. Friedman, L. M. Higgins, and J. Gore, Nature Ecology & Evolution **1**, 0109 (2017), ISSN 2397-334X, URL http://www.nature.com/articles/s41559-017-0109.
[S16] R. L. Clark, B. M. Connors, D. M. Stevenson, S. E. Hromada, J. J. Hamilton, D. Amador-Noguez, and O. S. Venturelli, Nature Communications **12**, 3254 (2021), ISSN 2041-1723, URL http://www.nature.com/articles/s41467-021-22938-y.
[S17] R. L. Clark, B. M. Connors, D. M. Stevenson, S. E. Hromada, J. J. Hamilton, D. Amador-Noguez, and O. S. Venturelli, GitHub repository (2021), https://github.com/RyanLincolnClark/DesignSyntheticGutMicrobiomeAssemblyFunction.
[S18] B. Wohlberg, Proceedings of the 16th Python in Science Conference pp. 1–8 (2017), conference Name: Proceedings of the 16th Python in Science Conference, URL https://conference.scipy.org/proceedings/scipy2017/brendt_wohlberg.html.
[S19] S. S. Chen, D. L. Donoho, and M. A. Saunders, SIAM Journal on Scientific Computing (2006), URL https://epubs.siam.org/doi/10.1137/S1064827596304010.
[S20] T. Hastie, R. Tibshirani, and M. Wainwright, *Statistical Learning with Sparsity* (Routledge, Boca Raton, 2015), 1st ed., ISBN 978-1-4987-1216-3.
